# Patient-Specific Vascularized Lung Tumor Organoids for Tumor-Immune Profiling

**DOI:** 10.64898/2026.06.01.729448

**Authors:** Naveen R. Natesh, Surjendu Maity, Raina Kikani, Sajeeshkumar Madhurakkat Perikamana, Giwon Cho, Nicole Angel, Zhicheng Ji, Shyni Varghese

## Abstract

The use of cellular systems to advance cancer therapeutics has expanded rapidly, spanning cell therapies to patient-specific tumor models. Platforms that recapitulate key features of the tumor microenvironment, including vascular and immune components, hold significant potential to improve the predictive power and translational relevance of preclinical models. Here, we report a vascularized tumor organoid platform that combines self-organizing microvascular networks with patient-derived tumor organoids and tumor-infiltrating lymphocytes. To minimize non-specific endothelial immunogenicity and enable broader compatibility across patient samples, we engineered the vasculature using β2-microglobulin-knockout endothelial cells. Leveraging this system, we established patient-specific, lymphocyte-incorporated tumor models that enabled quantitative assessment of T cell infiltration. In conjunction with immune checkpoint blockade, this platform distinguishes responder and non-responder patient samples, consistent with the clinical observations. Single-cell RNA-sequencing revealed tumor-intrinsic and immune-associated programs underlying this stratification, identifying tumor-driven hyperangiogenic signaling as a barrier to T cell extravasation. Pharmacological co-targeting of PD1 and VEGF restored T cell infiltration in non-responder organoids, shifting them from an immune-excluded to an immune-inflamed state. Together, this vascularized tumor organoid platform provides a predictive and mechanistic framework for modeling patient-specific immunotherapy responses and design of combination therapies.

## INTRODUCTION

Immunotherapies have emerged as a major modality in cancer treatment, transforming the therapeutic paradigm and offering durable clinical responses in a subset of patients. A canonical example is immune checkpoint blockade targeting programmed cell death protein 1 (PD-1), programmed death-ligand 1 (PD-L1), or cytotoxic T-lymphocyte–associated protein 4 (CTLA-4), which is now standard of care for multiple cancer types [1]. Cellular immunotherapies like chimeric antigen receptor (CAR) T cells and tumor-infiltrating lymphocytes (TIL) have also been demonstrated as an effective treatment for several blood cancers and PD1-refractory metastatic melanoma [2]. Despite these advances, extending the benefits of immunotherapy to solid tumors, which account for most of the cancer-related mortality, remains a major challenge. Cellular immunotherapies exhibit modest efficacy in solid tumors, owing in part to intratumoral heterogeneity, variable antigen presentation, and the complex tumor microenvironment, including spatial gradients of soluble mediators and chemokines that regulate immune cell trafficking and function [3–6]. Experimental models that recapitulate, even partially, key features of the tumor organ are therefore critical for evaluating efficacy of immunotherapies. Furthermore, such models can be used to gain a deeper understanding of the key aspects of tumor-immune interactions and the mechanisms by which certain tumors subjugate the immune system, ultimately improving therapeutic outcomes. Although syngeneic and transgenic mouse models have been instrumental in preclinical research, they are constrained by species-specific differences, limited patient specificity, and practical considerations [7–10]. Hence, there has been a surge of interest in developing complementary, human-relevant platforms for mechanistic interrogation and therapeutic testing.

Microengineered organotypic models, such as spheroids, organoids, assembloids, and explants recapitulate key structural and functional features of tumors and disease states [11–17]. Integration of these systems with microfluidic technologies has enabled the development of three-dimensional (3D) tumor(oid)-on-chip platforms that incorporate multicellular architecture, barrier interfaces, controlled microenvironments, and even perfusable vasculature. These microphysiological platforms offer unique opportunities to model key hallmarks of tumor, including cellular heterogeneity and dynamic cell-cell interactions, and have increasingly been applied to study patient-specific drug responses [14, 18–21]. More recently, such platforms have been extended towards understanding tumor-immune interactions [22–24]. Employing vascularized tissue models for cell therapy testing requires a perfusable system with cellular units uniquely infiltrated by or adjacent to capillary networks. Yet, current platforms are scarce and largely rely on endothelial monolayers functioning as a ‘capillary endothelium wall’ that can be transmigrated by immune cells or heterogenous tumor explants embedded within vascular networks.

Inspired by the promise of vascularized tumor-on-chip platforms, we developed a patient-specific vascularized tumor-on-chip platform for preclinical evaluation of drug responses and cell–based immunotherapies. This system integrates the perfusable microvascular network with patient-derived tumor organoids, enabling continuous exposure of immune cells and therapeutics through a physiologically relevant vascular interface. Establishing such multicellular systems presents key challenges, including the coordination of distinct cellular populations with divergent growth and maturation requirements, as well as the maintenance of long-term functional stability. In addition, endothelial immunogenicity can introduce confounding signals that obscure interpretation of immune-mediated tumor responses.

To enhance platform robustness and generalizability, we engineered the vasculature using β*2*-microglobulin (B2M)-deficient endothelial cells, generated via CRISPR–Cas9 gene editing to minimize allogenic immune recognition. Using patient-derived adenocarcinoma organoids and autologous TILs, we evaluated T cell extravasation toward the tumor compartment in the presence of immune checkpoint blockade. Retrospective comparison with the patient’s baseline tumor characteristics showed that the platform recapitulated key patient-specific features observed clinically. Furthermore, single-cell RNA sequencing, combined with pharmacological perturbation, revealed tumor- and immune-intrinsic programs associated with therapeutic response, including a shift from an immune-excluded to an immune-inflamed state driven by enhanced T cell recruitment.

## RESULTS

### Multi-cellular lung tumor organoid with perfusable vasculature

To generate a perfusable, vascularized patient-derived lung tumor organoid (LTO) model, we developed a multicompartment microfluidic device that supports spatially confined culture of multiple cell types while enabling intercellular interactions. The device is comprised of four compartments, as shown in the schematic (**Fig. 1a**). A detailed schematic with dimensions is provided in **Fig. S1a,** and the experimental workflow illustrated in **Figures 1b and S1b**. The multicompartment, modular design of the device with designated compartments for the localization of different cell populations, including organoids, makes the platform highly versatile. The device can be readily adapted by altering compartment size, geometry, or number to accommodate additional cell types or experimental conditions. Different tumor organoids, explants, or microtissues can be integrated with flexible timing of transplantation. The current device configuration includes a microvascular network (mVN) compartment (i), a central circular chamber for the LTO (ii), side chambers for fibroblasts (iii), and media channels (iv), separated by arrays of micropillars that enable hydrogel entrapment and fluid flow (**Fig. 1a**). Fibroblasts, in addition to being a key stromal cell population, secrete factors that promote microvascular network formation and maturation [25].

**Figure 1:**
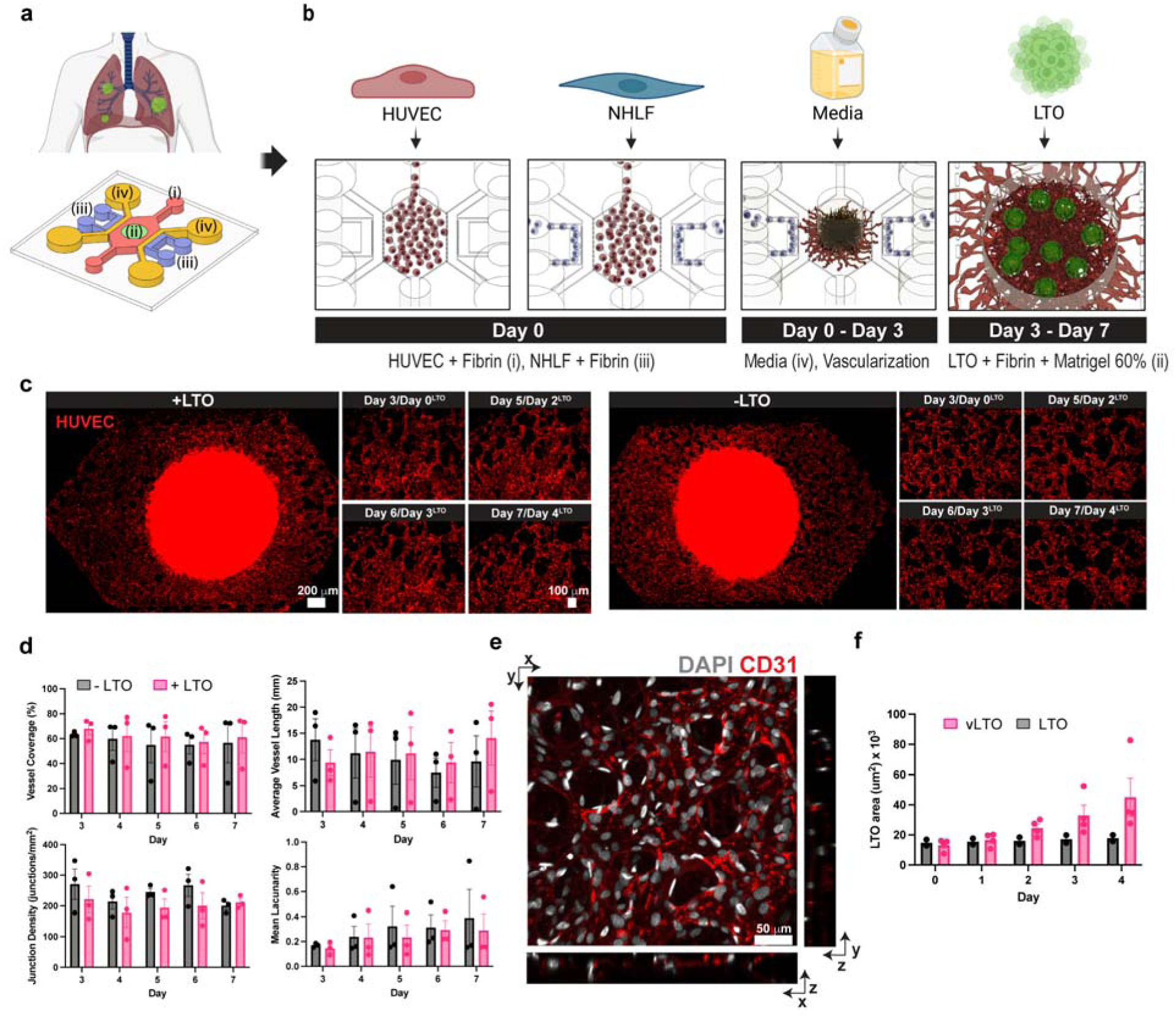
Vascularized lung tumor platform. (a-b) Conceptual illustration and schematic of the device. Microvascular compartment (i), organoid compartment (ii), fibroblast compartment (iii), and media channels (iv). (b) Timeline of cell seeding including LTO in the device: On day 0, devices are seeded with HUVEC and NHLF encapsulated in a fibrin hydrogel; devices are cultured for 3 days prior to LTO transplantation; devices are then cultured for an additional 4 days before characterization. (c) Representative images of the vascularized LTO platform with or without LTO incorporation. (d) Quantification of microvasculature network morphology metrics with or without LTO implantation; N = 3 independent devices; data are presented as mean +/−SEM; Repeated measures (RM) two-way ANOVA with Šídák’s multiple comparisons test yielded no significant differences in any of the metrics (p > 0.05). (e) Confocal microscopy images and z-stack sectioning of microvessels in the vascularized LTO platform. (f) Quantification of LTO area in static culture versus the vascularized LTO platform; N(LTO) = 2 wells; n(LTO) = 8 LTO/well, N(vLTO) = 4 devices, n(vLTO) ≥ 2 vLTO/device; data presented as mean +/− SEM; RM two-way ANOVA with Šídák’s multiple comparisons test yielded no significant differences (p > 0.05).

Human umbilical vein endothelial cell (HUVEC)-laden fibrin gels in compartment *i* self-organized into perfusable microvascular networks by day 3 post-seeding, as confirmed by confocal imaging and quantitative network analysis (**Fig. 1c-e, S1c**). Network morphology was assessed over time using AngioTool, a semi-automated software widely employed to analyze vascular network both *in vitro* and *in vivo* [26]. Consistent with previous reports, the microvascular networks exhibited high vessel coverage and connectivity, reflected by increased branch-point frequency, low lacunarity (a measure of structural uniformity), and stable mean vessel length (**Fig. 1d**) [27].

We next examined the effect of incorporating lung tumor organoids on microvascular network formation. Patient-derived lung adenocarcinoma organoids were generated as described in the Methods. Lung tumor organoid (LTO)-laden fibrin/Matrigel hydrogel was introduced into compartment *ii* of the device on day 3 following incorporation of HUVECs and NHLFs. Day 3 was chosen because the HUVECs had reproducibly self-organized into a stable microvascular network by this time point. The multi-cellular culture was maintained in a 1:1 mixed medium consisting of EGM2 MV and LTO medium. The medium composition and matrix components were optimized to support the different cell populations in the coculture, including the LTOs. Characterization of the microvascular network following the LTO incorporation revealed vessel characteristics comparable to those observed in the absence of tumor organoids, suggesting that the LTO incorporation did not significantly alter the microvascular network formation or morphology (**Fig. 1d**). As expected, the presence of a perfusable microvascular network enhanced organoid growth, with vascularized LTOs exhibiting larger size compared to non-vascularized controls (**Fig. 1f, S1d**).

Confocal imaging revealed that the vascular networks interfaced with the LTOs, with endothelial sprouts extending towards and into the organoids (**Fig. 2a**). Owing to the relatively large size and dense three-dimensional (3D) architecture of the organoids embedded within the matrix and associated microvascular network, confocal imaging in combination with IMARIS software was used to visualize and quantify the vascular–LTO interface using 3D rendering and distance transformation analysis (**Fig. S2**). Spatial distance distribution analysis of the micro vessel–tumor organoid interface was performed for multiple organoids to quantify the extent of vessel contact. A modal value of approximately 0 μm in the frequency distribution indicates that most vessels are in direct contact with the LTOs. Negative values (<0 μm) suggest vessels penetrating into the organoid, whereas positive values (>0 μm) indicate vessels located outside the organoid boundary. These analyses indicate that most LTOs exhibit a high frequency of close vessel–organoid contact, with extensive vascular integration observed across multiple tumor organoids (**Fig. 2b**). Quantification of the spatial distribution indicated an average contact surface area of approximately 40 μm^2^ between the tumor organoids and the microvascular vessels (**Fig. 2c**).

**Figure 2:**
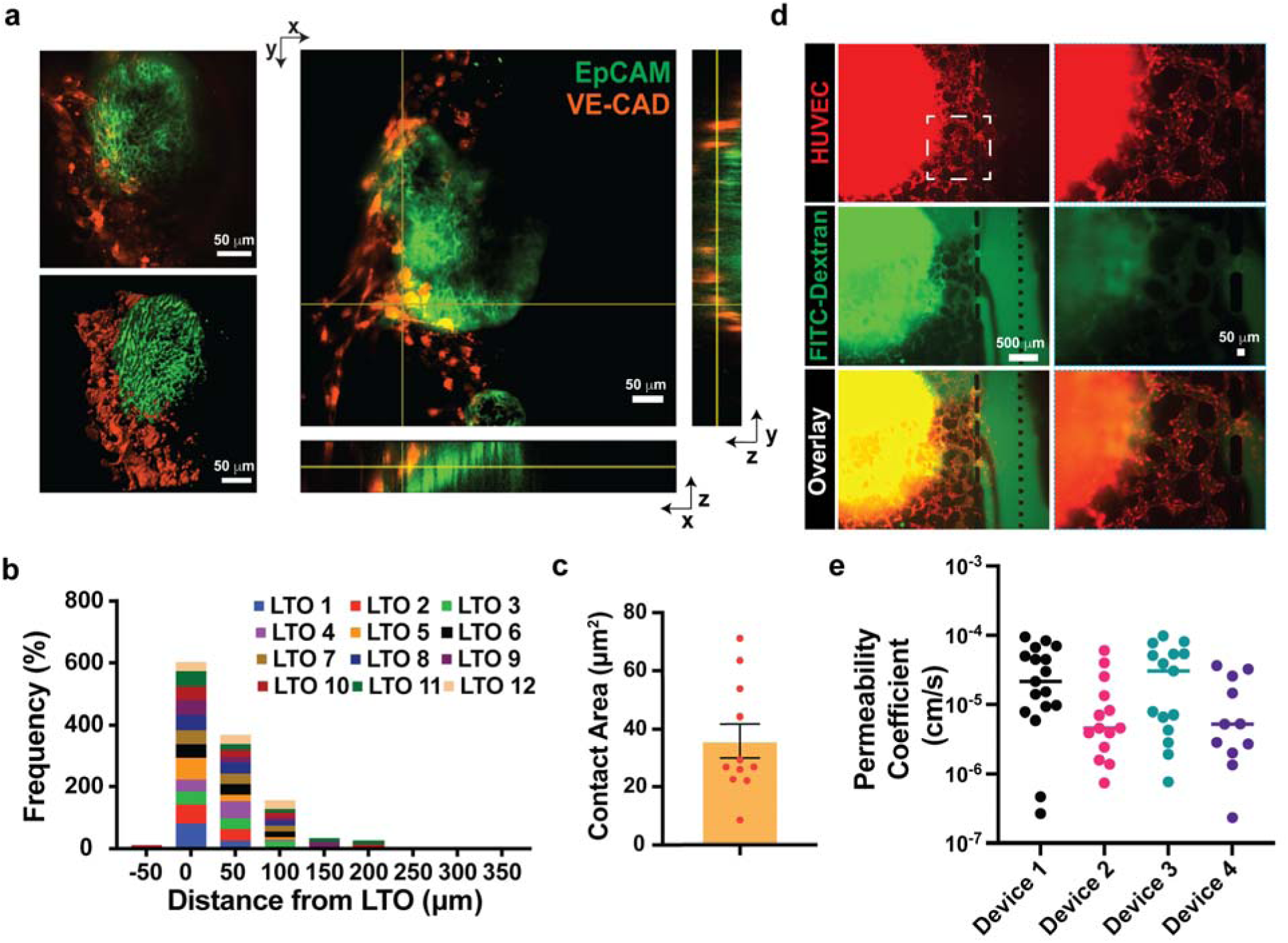
Characterization of lung tumor organoid vascularization within the platform. (a) Representative confocal microscopy images of vascularized lung tumor organoid with z-stack sections. (b) Frequency distribution of vascular-LTO contact distances. (c) Contact area between vascular endothelial cells and LTO; N = 2 devices, n ≥ 4 vLTO/device; data are presented as mean +/− SEM. (d) Representative image of FITC-Dextran perfusion in the vascularized LTO platform. (e) Quantification of permeability coefficients across multiple devices using FITC-dextran perfusion; N = 4 devices; n ≥ 11 microvessels per device; Brown-Forsythe ANOVA with Dunnett’s T3 multiple comparisons test yielded no significant differences (p > 0.05).

Perfusability of the microvascular network in the presence of LTO was confirmed by perfusion with 2000kDa FITC-Dextran. The fluorescent tracer was introduced through the media inlets into the microvascular channels, which remained confined within the vascular lumen while reaching the tumor compartment (**Video S1 and S2, Fig. 2d**). Vessel permeability measurements indicated the formation of a robust and relatively low-permeability vascular network, with coefficients comparable to those reported for engineered microvascular systems (**Fig. 2e**) [28, 29].

### Vascularized tumor organoid models to assess drug toxicity

We used the vascularized LTO model to evaluate drug toxicity using cisplatin as a model chemotherapeutic agent. Cisplatin is a first-line therapy used in the treatment of non–small-cell lung carcinoma [30]. The drug was introduced via perfusion through the microvascular network and exposing the tumor organoids to varying concentrations of cisplatin. In parallel, non-vascularized lung tumor organoids were exposed to the same drug concentrations. Both vascularized and non-vascularized tumor organoids exhibited comparable responses to cisplatin, albeit with subtle differences (**Fig. 3a**). Cell viability measurements at 24 hours post-treatment showed a concentration-dependent cytotoxic effect, with significant cell death observed in both culture conditions at cisplatin concentrations of 10 μM or higher (**Fig. 3b, c**). Notably, we observed a striking cell type-specific cytotoxic response in the vascularized tumor organoid model, wherein the peripheral microvascular cells exhibited minimal toxicity following cisplatin treatment (**Fig. 3d, e**). Intrigued by this finding, we further examined the effect of cisplatin on HUVECs in conventional monolayer culture, which showed significant cell death at 100 μM concentration (**Fig. S3**).

**Figure 3:**
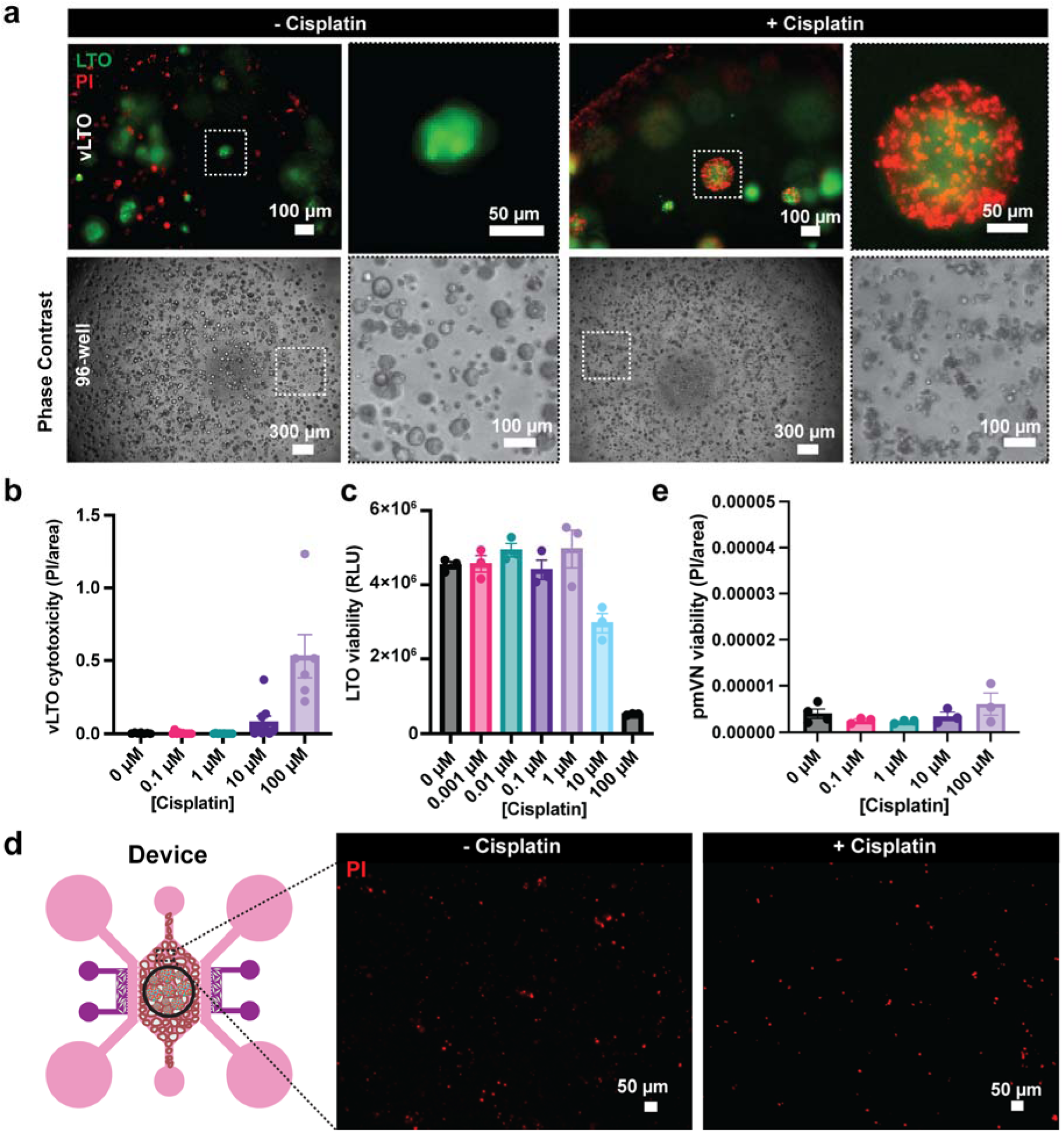
Drug response testing using the vascularized tumor organoid platform. (a) Representative images from vascularized LTOs and non-vascularized LTOs treated with cisplatin. (b) Quantification of LTO cytotoxicity upon cisplatin treatment of varying concentrations in the vascularized LTO platform; N(vLTO) = 2 devices, n(vLTO) ≥ 6 vLTO/device; data presented as mean +/− SEM. (c) Quantification of LTO viability in non-vascularized LTO following cisplatin treatment with varying concentrations; N(LTO) = 3 wells; data presented as mean +/− SEM. (d) Representative images of peripheral microvasculature within the vascularized LTO device with or without cisplatin treatment. (e) Quantification of cytotoxicity of peripheral microvasculature following cisplatin treatment of varying concentrations; N ≥ 3 devices per group.

### Vascularized tumor organoid models to assess T cell-tumor interactions

We employed the vascularized LTO model to investigate immune cell extravasation and infiltration into tumor organoids and the surrounding microenvironment. Building on our previous study, we first examined the effect of THP-1 monocytes in facilitating CD8⁺ T cell recruitment into tumor organoids [24]. Sequential perfusion of cells was performed, wherein the monocytes were first introduced into the microvascular network, followed 2 hours later by perfusion of CD8⁺ T cells for 24 hours (**Fig. 4a**). Confocal imaging revealed a significantly higher accumulation of infiltrating CD8⁺ T cells in vascularized LTOs in the presence of monocytes compared to their absence (**Fig. 4b-d**). Spatial distribution analysis of CD8⁺ T cell localization within tumor organoids and its immediate surrounding demonstrated significantly increased T cell recruitment into the organoid compartment when monocytes were present, as shown by the CD8⁺ T cell frequency distribution (**Fig. 4d**). Quantitative analysis of the frequency distribution further showed a higher density of CD8⁺ T cells within 100 μm of the LTOs in the presence of monocytes across multiple vascularized LTOs (**Fig. 4c**).

**Figure 4:**
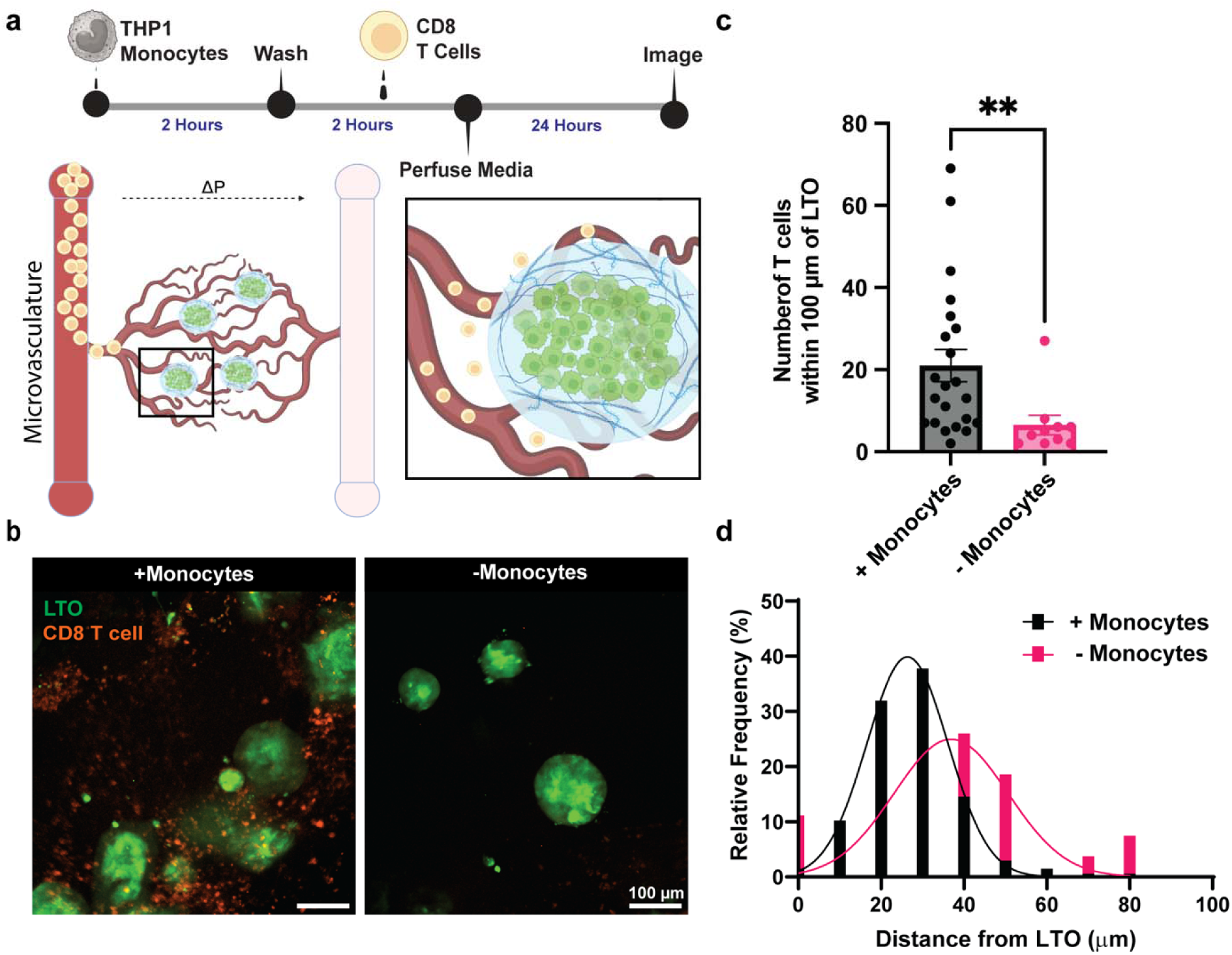
T cell recruitment using vascularized LTO platform. (a) Schematic illustrating the experimental workflow: Incorporation of THP-1 monocytes followed by perfusion of primary CD8^+^ T cells through the microvascular network. (b) Representative confocal microscopy images of vascularized LTO infiltrated by T cells in the presence or absence of monocytes. (c) Quantification of T cells localized near the vLTO in presence or absence of monocytes (N = 3 devices per group, n ≥ 2 vLTO/device). Data are presented as mean +/− SEM; Unpaired t-test with Welch’s correction; ** = p < 0.005. (d) Frequency distribution of the distance between T cells and vLTO in the presence or absence of monocytes (Gaussian fit shown in black and pink curves, respectively; N = 3 device per group, n ≥ 2 vLTO/device).

### Towards a generalized microvascular platform for tumor–immune interactions

The use of primary human endothelial cells in T cell recruitment assays may introduce bias or non-specific T cell activation due to recognition of allogenic major histocompatibility complex class I. β2-microglobulin (B2M) is an essential component of the functional MHC-1 complex, thus genetic ablation of *B2M* represents a strategy to minimize artificial T cell activation by endothelial cells within the microvascular system [31, 32]. To this end, we employed CRISPR-Cas9-mediated gene editing to knock out *B2M* in HUVECs. A Cas9 ribonucleoprotein complex containing a single guide RNA (sgRNA) targeting exon 1 of the *B2M* locus was electroporated into HUVECs. The efficiency of the *B2M* knock-out was confirmed by flow cytometry, followed by sorting and expansion of *B2M*-negative (*B2M*-KO) HUVECs for cryopreservation and downstream studies (**Fig. S4a**). Tracking of Indels by DEcomposition (TIDE) analysis of genomic DNA from *B2M*-KO HUVECs confirmed the presence of insertions and deletions at the CRISPR target site (**Fig. S4b**) [33]. Mutations within the sgRNA-targeted region were further validated by Sanger sequencing (**Fig. S4c**). We next evaluated the ability of *B2M*-KO HUVECs to self-organize into perfusable microvascular networks within the microfluidic device. Fluorescence microscopy and quantitative analyses demonstrated that *B2M*-KO HUVECs formed perfusable networks with vessel parameters comparable to those formed by unedited HUVECs (**Fig. S5a**). To evaluate immunogenicity, we performed co-culture experiments using CD8⁺ T cells with either *B2M*-KO HUVECS or unedited HUVECs and quantified interferon-γ (IFN-γ) secretion as a readout of T cell activation. ELISA measurements showed a ∼5-fold decrease in IFN-γ levels in co-cultures with *B2M*-KO HUVECs compared to those with unedited HUVECs (**Fig. S5b**). Together, these findings suggest the successful establishment of *B2M*-KO HUVECs and their ability to form perfusable microvascular networks with attenuated allogeneic immune responses.

### Vascularized patient-matched tumor organoid platform for tumor-immune interactions

To fully establish a patient-agnostic vascularized platform, we created vascularized tumor organoid platform using *B2M*-KO HUVECs and investigated its applicability for testing immune checkpoint blockade with patient-derived tumor-infiltrating lymphocytes (TILs). Patient-matched TILs were collected from patients with lung adenocarcinoma, expanded *in vitro*, and subsequently used for the studies (**Fig. 5a**). LTOs were also generated from the corresponding patient tumor samples, and H&E staining revealed cellular morphology and structural integrity of the patient-derived tumor organoids (**Fig. 5c**). The LTOs were further characterized by immunofluorescent staining for lung adenocarcinoma markers such as CK7, Napsin, and EpCAM (**Fig. 5d**). Representative H&E images of the patient tumor samples are shown in **Supplementary Figures S6a and S6b**. As outlined in the experimental workflow in **Fig. 5b**, the LTOs were incorporated into the vascularized device, followed by sequential perfusion of the microvascular networks with medium containing THP-1 monocytes, patient matched TILs, and subsequently medium supplemented with anti-PD1 to assess the impact of immune checkpoint inhibition on TIL recruitment.

**Figure 5:**
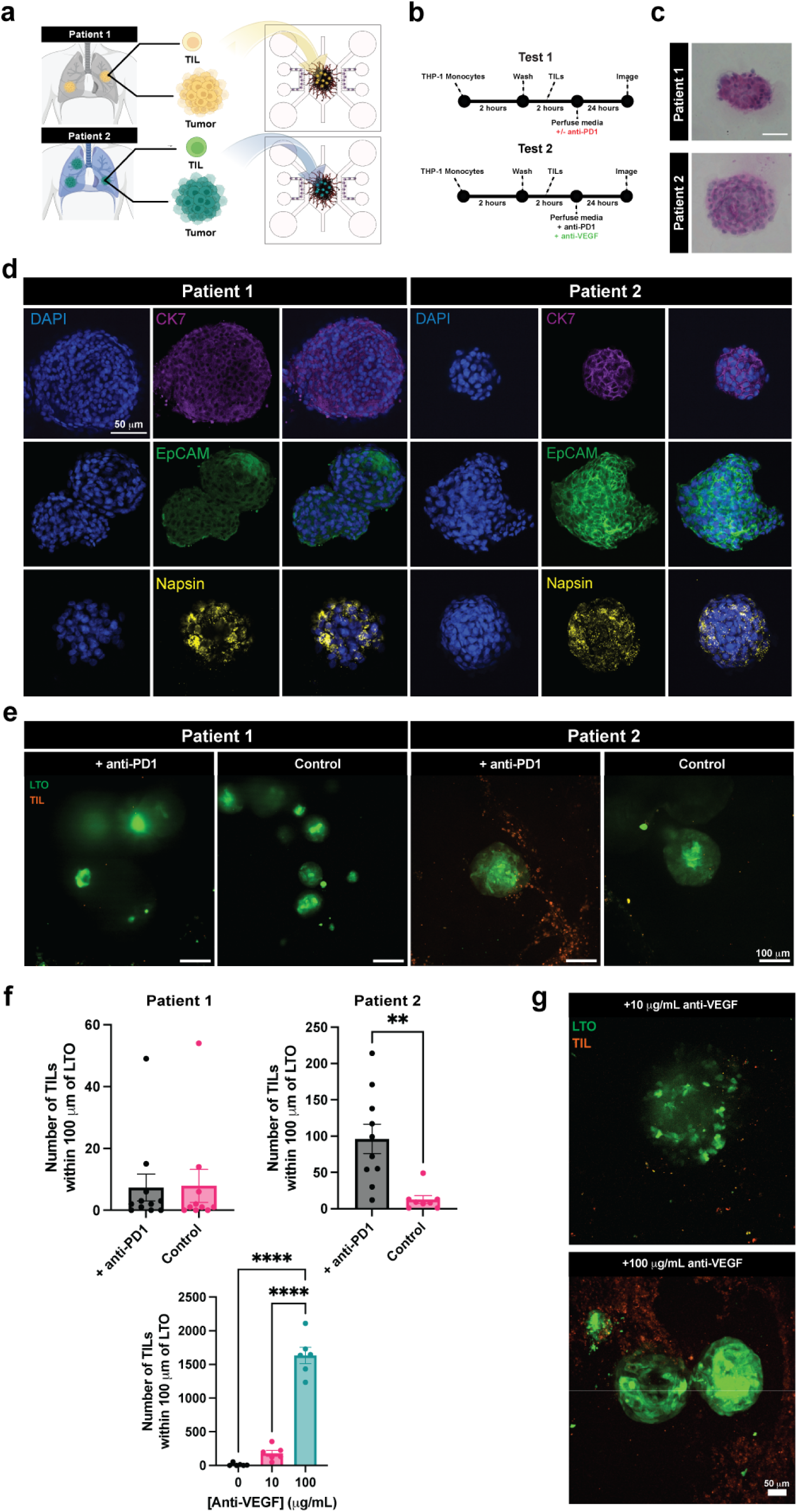
Patient-specific vascularized LTO platform for autologous TIL recruitment. (a) Schematic illustrating patient sample processing and generation of LTOs and matched TILs. (b) Schematic of perfusion and drug treatment workflow. Incorporation of THP-1 monocytes followed by perfusion of patient matched TILs through the microvascular network in the presence of anti-PD1 or anti-VEGF antibody. (c) Hematoxylin and eosin (H&E) staining of LTO generated from patients with lung adenocarcinoma. (d) Immunofluorescence staining of patient-derived LTOs for different lung adenocarcinoma markers; blue: DAPI, magenta:CK7, green:EpCAM, yellow: Napsin. (e) Representative confocal microscopy images of vascularized LTOs from patient 1 and 2 in the presence or absence of anti-PD 1 treatment. (f) Quantification of T cells localized near vLTO in the presence or absence of anti-PD1 (N ≥ 2 devices per group, n ≥ 8 vLTO/device. Data are presented as mean +/− SEM; unpaired t-test with Welch’s correction; ** = p < 0.005) and in dual anti-VEGF/anti-PD1 blockade (N ≥ 2 devices per group, n ≥ 1 vLTO/device. Data are presented as mean +/− SEM; Ordinary one-way ANOVA with Tukey’s multiple comparisons test; * = p < 0.05, **** = p < 0.0001). (g) Representative confocal microscopy images of Patient 1 vLTO in the presence of both anti-PD1 and anti-VEGF antibody treatment.

Confocal imaging of the tumor organoids treated with anti-PD1 revealed patient-specific differences in TIL extravasation and infiltration compared with untreated controls (i.e., not exposed to anti-PD1) (**Fig. 5e**). Moreover, TILs from Patient 1 showed no significant recruitment following anti-PD1 treatment, with few TILs detected within the LTO or within 100 μm of its periphery (**Fig. 5e**). In contrast, TILs from Patient 2 extravasated into the tumor organoid, with abundant presence within the organoid and its immediate vicinity in response to anti-PD1 treatment (**Fig. 5e, f**). These findings suggest that enhanced TIL infiltration and interaction with tumor cells in this platform may represent a patient-specific response to immune checkpoint blockade, likely influenced by intrinsic differences in tumor characteristics and/or TIL functional states. We next assessed how these observations compared with patient clinical data, which noted that Patient 2’s tumor had a median of five intraepithelial lymphocytes (IELs) per 10 tumor cells, whereas Patient 1 had fewer than one IEL per 10 tumor cells (**Supplementary Table S1**). In addition, immunofluorescence analysis of biopsied patient tumor tissue samples demonstrated significantly more CD3^+^ cells in Patient 2 compared to Patient 1, indicating a higher abundance of intratumoral T cells in Patient 2 (**Fig. S6c, d**). Given that intratumoral lymphocyte abundance can predict response to immune checkpoint blockade, the vascularized LTO platform independently recapitulated patient sensitivity to anti-PD1 therapy [34–36].

### Single-cell sequencing to determine the patient response

To understand the mechanisms underlying the observed functional readouts, we performed single-cell RNA sequencing (scRNA-seq) analysis of organoid platform containing lung tumor organoids with the immune cells. A total of 25,451 cells were sequenced and visualized using uniform manifold approximation and projection (UMAP) (**Fig. 6a**). Nine distinct cell types were annotated based on canonical marker expression (**Fig. 6a, b**; **Fig. S7a-d**). Patient 1 samples contained a higher proportion of tumor cells, possibly reflecting reduced T cell-mediated cytotoxicity, whereas the relative frequencies of other cell types were largely comparable between the two patients (**Fig. 6c**). Differential gene expression analysis identified several differentially expressed genes (DEGs) across cell types (**Fig. S7e-j**), with the most notable changes in elevated *VEGF* expression in LTOs and increased expression of *GZMB* and *PDL1* in CD8^+^ cytotoxic T cells (**Figure 6d-f**). Gene Ontology (GO) analysis revealed that genes upregulated in Patient 1 LTOs were enriched for pathways associated with positive regulation of angiogenesis and endothelial cell migration (**Fig. 6g**). In contrast, genes upregulated in Patient 2 LTOs were enriched for pathways related to T cell receptor signaling and activation (**Fig. 6h**). Notably, LTOs from Patient 1 exhibited marked upregulation of *VEGF*, which may contribute to the attenuated response of anti-PD1 therapy, potentially through enhanced PD-L1 expression in CD8^+^ T cells (**Fig. 6i, j**) Additionally, T cells from Patient 1 displayed higher median exhaustion scores compared to those from Patient 2 (**Fig. 6k**). Together, these data suggest the contribution of both tumor- and TIL-intrinsic mechanisms for the reduced T cell infiltration observed in Patient 1 (**Fig. 6l**).

**Figure 6:**
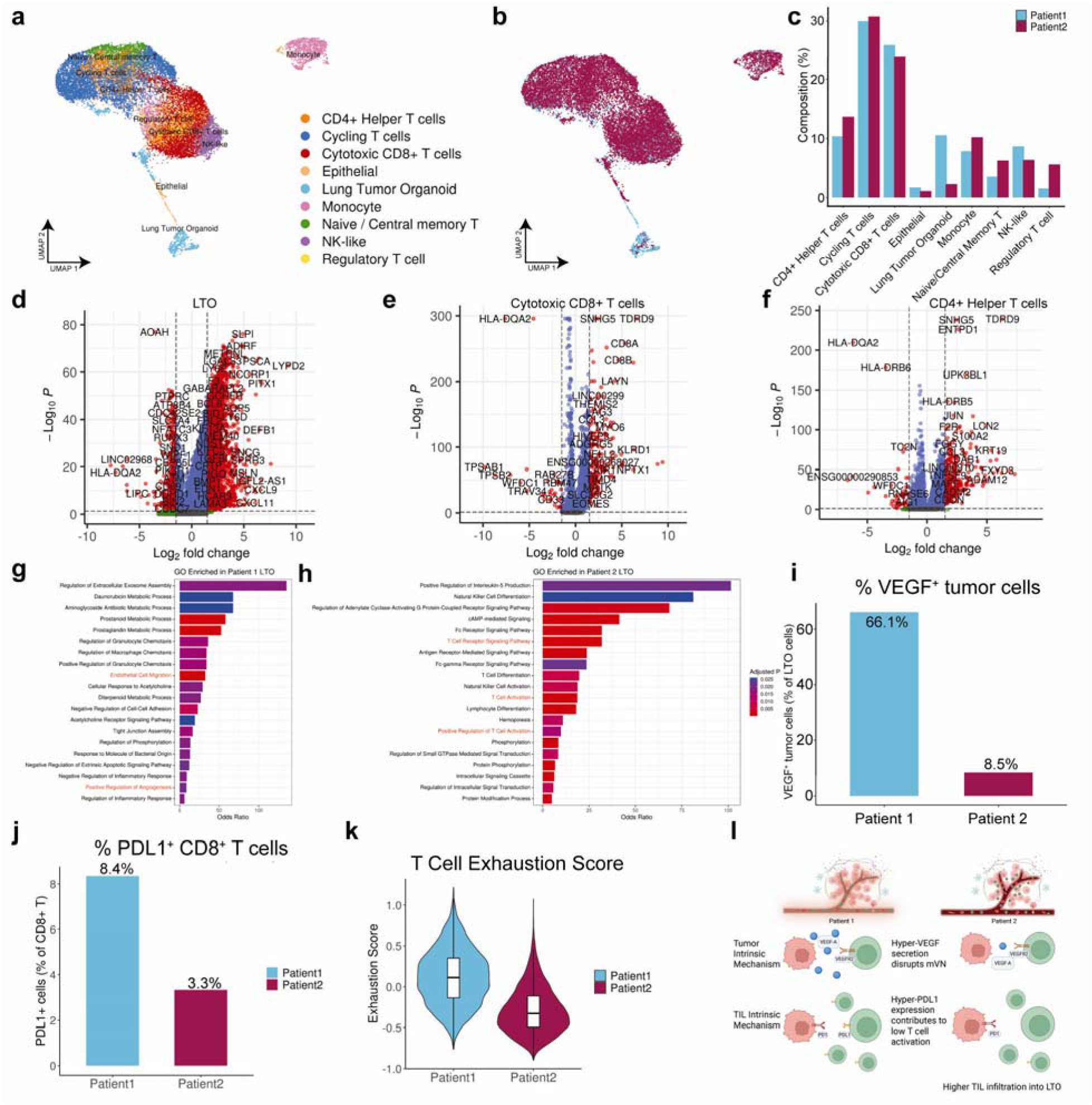
Single-cell RNA sequencing analysis of LTOs. (a) UMAP visualization colored by cell type. (b) UMAP visualization colored by patient. (c) Cell type composition across patients. (d-f) Differentially expressed genes in LTO cells, CD8^+^ T cells, and CD4^+^ helper T cells respectively (|Log2 fold change| ≥ 1.5, adjusted p value < 0.05). (g-h) Top 20 enriched Gene Ontology biological processes for Patient 1 and Patient 2, respectively (adjusted p value < 0.05). (i) Percentage of VEGF^+^ LTO cells per patient. (j) Percentage of PD-L1^+^ CD8^+^ T cells per patient. (k) T cell exhaustion score per patient. (l) Proposed mechanism illustrating tumor-intrinsic and TIL-intrinsic mechanisms that limit T cell recruitment to and infiltration into LTOs.

### Dual antibody treatment of vascularized LTO to promote TIL trafficking

Vascular endothelial growth factor (VEGF) is a pleiotropic growth factor that plays a central role in regulating angiogenesis under both homeostatic and pathological conditions. The proangiogenic molecule, VEGF, secreted by tumor cells, has been recognized as a key mediator of tumor immune evasion. The PD1-inducing effects of hyperactive VEGF signaling have also been well established [37]. Based on these findings, we reasoned that although T cells may home to regions of the vascular networks adjacent to lung tumor organoids, their extravasation could be impeded by tumor-associated signaling that contributes to immunosuppressive pathways. We therefore examined whether dual blockade of PD1 and VEGF could restore T cell extravasation and localization to LTOs derived from Patient 1. Using the vascularized LTO platform, we assessed TIL recruitment and extravasation in response to anti-PD1 in the presence of an anti-VEGF antibody (**Fig. 5b**). Treatment of LTOs with anti-VEGF and anti-PD1 resulted in a significant enhancement of T cell recruitment (**Fig. 5g**). Specifically, the number of T cells within the vascularized LTOs and in their immediate vicinity was significantly increased in the presence of anti-VEGF compared to controls not treated with the VEGF antibody. T-cell infiltration increased in a dose-dependent manner with increasing concentrations of the VEGF antibody (**Fig. 5f, g**). Collectively, these results suggest that high levels of lung tumor cell-secreted VEGF can modulate endothelial cells of the microvascular network and contribute to impaired TIL extravasation and infiltration, and that VEGF blockade can at least partially alleviate this immunosuppressive barrier.

## DISCUSSION

The development of human relevant preclinical models has gained significant attention in recent years, particularly in light of the FDA Modernization Act 2.0 and the Complement Animal Research in Experimentation (Complement-ARIE) program, which encourage the adoption of alternative models for evaluating therapeutic efficacy. Among new approach methodologies (NAMs), advanced cellular systems have emerged as an important tool. Although substantial progress has been made with three-dimensional (3D) models such as organoids and spheroids, there is growing recognition of the importance of incorporating vasculature to better recapitulate tissue physiology. Vascularization not only enhances nutrient and oxygen delivery but also introduces critical components of the tissue microenvironment. In diseases such as cancer, the vasculature plays a central role in tumor progression and therapeutic outcomes, including responses to immunotherapy. The vascularized tumor organoid platform described here could be a technological advancement in the field.

Although, like other organ-on-chip platforms, our system does not incorporate all components of the complex tumor microenvironment, the platform incorporates the minimal set of components necessary to address the biological question(s) under investigation. A key advantage of this platform is its modular design, which enables systematic and controlled interrogation of defined variables. The platform supports seamless tumor integration while allowing independent manipulation and recombination of various cellular components within designated compartments. This approach facilitates mechanistic dissection of tumor–microenvironment and their influence on therapeutic outcomes. Moreover, the strategy of independently establishing individual components prior to assembly mitigates common challenges associated with complex co-culture systems, including medium incompatibility, matrix selection constraints, adverse cell–cell interactions, and disproportionate overgrowth of specific cellular units. In the present study, normal lung fibroblasts were incorporated in place of cancer-associated fibroblasts to enable a focused investigation of tumor–immune interactions while minimizing confounding stromal effects.

Our findings highlight the functional importance of vascularization. While the incorporation of lung tumor organoids (LTOs) had minimal impact on microvascular network formation, the presence of a perfusable vascular network significantly enhanced organoid growth and viability, likely through improved nutrient delivery and waste removal. In drug response studies, the chemotherapeutic agent cisplatin induced comparable cytotoxic effects in both vascularized and non-vascularized LTOs. Notably, however, endothelial cells within the peripheral 3D vascular network exhibited minimal sensitivity to cisplatin across the tested concentrations, in contrast to endothelial monolayer cultures, which displayed high toxicity. These differences in drug response may be attributed to the fundamental differences between 2D and 3D cultures, with the latter more closely recapitulating physiological architecture and barrier function.

When adapting vascularized platforms for the evaluation of cell-based immunotherapies, the immunogenicity of the vascular component is a key challenge. To address this, we engineered microvascular systems using β2-microglobulin (B2M) knockout endothelial cells generated through genome engineering. The generation of *B2M*-KO HUVECs-derived microvascular network is an important step forward in developing patient-specific models to study cell therapies. Employing these engineered endothelial cells in tandem with patient-matched LTOs and TILs enabled the assessment of patient-specific immune responses, including differential sensitivity to PD1 blockade.

Single-cell RNA sequencing provided mechanistic insight, revealing tumor-intrinsic and immune-associated programs associated with the differential TIL response. In particular, hyperangiogenic signaling emerged as a key feature of non-responsive tumors. Elevated VEGF secretion in non-responder organoids suggests a plausible mechanism by which tumor-driven vascular signaling contributes to immune evasion. VEGF-mediated effects on endothelial cells, including induction of PD-L1 expression, may promote T cell dysfunction independently of canonical immune checkpoint pathways [37]. Moreover, excessive pro-angiogenic signaling is known to drive the formation of abnormal, leaky, and disorganized vasculature, which can impede both drug delivery and immune cell recruitment by creating physical and biochemical barriers [38–42]. Importantly, the dual inhibition of PD1 and VEGF signaling restored T cell extravasation and infiltration in non-responder organoids, further supporting hyperangiogenic signaling in observed immune exclusion. These findings align with clinical efforts exploring combinatorial or bispecific approaches of targeting both immune checkpoints and angiogenic pathways in solid tumors, particularly for non-small cell lung cancer [43–45]. Collectively, our results show that the vascularized tumor organoid platform described in this study can not only recapitulate patient-specific therapeutic responses but also enables the identification effective combination therapies in a controlled, mechanistic setting.

In conclusion, we developed a patient-agnostic, vascularized tumor organoid platform incorporating patient-derived cells for the evaluation of drug response and immune cell therapies. The platform captures patient-specific variability in T cell extravasation and responses to immune checkpoint blockade and provides mechanistic insight into resistance pathways. Furthermore, the platform enabled testing of combination therapies, as demonstrated by the restoration of T cell infiltration following dual PD1 and VEGF inhibition. With further validation across larger and more diverse patient cohorts, this platform has the potential to serve as a predictive and mechanistic framework for preclinical immunotherapy assessment. Given the central role of vascular dysfunction and hyperangiogenic signaling across solid tumors, the approach is broadly applicable beyond lung cancer. Moreover, the modular nature of the platform allows integration of additional model types and therapeutic modalities, including tumor explants, slice cultures, and engineered immune cell therapies such as CAR-T cells, thereby expanding its utility for next-generation cancer modeling and therapeutic development.

## MATERIALS AND METHODS

### Microfluidic device fabrication

The microfluidic device is composed of three distinct chambers designated to house different cell populations, including LTOs, and to support their respective functions (**Fig. 1a,b** and **Fig. S1a**). The device was fabricated using conventional soft lithographic techniques [46]. The fibroblast chamber measures 5 mm x 750 μm, the microvasculature chamber measures 7.4 mm x 4.7 mm, and the circular organoid chamber has a diameter of 3 mm. The medium inlet/outlet ports are 4 mm diameter, and the fibroblast seeding inlet/outlet ports are 2 mm in diameter. All microchannels are 100 μm in height. Briefly, SU-8 100 photoresist (Kayaku Advanced Materials) was spin-coated onto 100 mm silicon wafers. A 20,000 dpi film photomask with the device design (AutoCAD) was aligned and exposed to UV light at an intensity of 10 mW/cm^2^ for 55 s. The wafer was then developed in SU-8 developer for 10 min. Patterned wafers were silanized under vacuum with trichloro(1*H*,1*H*,2*H*,2*H*-perfluorooctyl) silane (MilliporeSigma, 448931) for 15 min at 25 °C. Polydimethylsiloxane (PDMS; Sylgard 184, Dow Corning) base and curing agent were mixed at a 10:1 (w/w) ratio, cast onto the SU-8 master, degassed under vacuum in a desiccating chamber. and cured at 60 °C for 3 h. The cured PDMS was peeled from the master mold, and access ports were created using biopsy punches of defined diameters: 4 mm for medium inlets and outlets, 2 mm for fibroblast chamber ports, 3 mm for the organoid chamber, and 1 mm for the central chamber inlet and outlet. Rectangular glass coverslips were clean with 70% ethanol and treated with oxygen plasma for 1 min prior to bonding to PDMS layers. Microfluidic channels were coated with 1 mg/mL poly-(D)-lysine (Sigma, P7886) and incubated for 4 h at 37 °C, rinsed thoroughly with DI water and dried at 60 °C to restore hydrophobicity. Devices were stored at room temperature and used within 72 hours of fabrication.

### Cell culture

Human umbilical vein-derived endothelial cells (HUVECs) were purchased from Lonza (CC-2519) and cultured in EGM-2MV medium (CC-3202). Normal human lung fibroblasts (NHLFs, Lonza, CC-2512) were cultured in FGM-2 medium (CC-3132). THP-1 monocytes were obtained from ATCC (TIB-202) and cultured in RPMI-1640 supplemented with 10% FBS and 0.05 mM 2-mercaptoehtanol (Gibco, 21985023). Human peripheral blood CD8^+^ T cells were purchased from STEMCELL Technologies (200-0164) and maintained in RPMI-1640 supplemented with 10% FBS and 1% Penicillin-Streptomycin (“T cell medium”; Sigma, P4333). CD8^+^ T cells were used initially to optimize the platform and evaluate T cell extravasation and infiltration to tumor organoids. Patient-derived LTOs and tumor-infiltrating lymphocytes (TILs) were cultured as described below. All cells were cultured at 37 °C in 5% CO_2_ and 21% O_2_.

### Patient sample processing and cell isolation

Tumor tissues were processed as previously described [47]. Biological samples were obtained from patients undergoing standard-of-care biopsy or surgical resection at Duke University Hospital under an approved Institutional Review Board (IRB) protocol (Pro00102781) for patients with lung adenocarcinoma. Briefly, freshly resected or biopsy-derived tumor specimens were mechanically minced into approximately 1–2 mm fragments and subjected to enzymatic dissociation in digestion medium consisting of 1 mg/mL collagenase (Sigma, 11088858001), 3 mM CaCl₂, 0.1 mg/mL DNase I (StemCell Technologies, 07900), 10 mM Y-27632 (StemCell Technologies, 72302), and 100 µg/mL Primocin (Fisher Scientific, NC9141851). Tissue fragments were incubated in digestion medium at 37 °C under gentle agitation for 30 min. If incomplete dissociation was observed, samples were further incubated for an additional 10–15 min until a single-cell suspension with minimal residual clumps was achieved. Following enzymatic digestion, cell suspensions were filtered through a 70 μm cell strainer to remove undigested material and aggregated tissue. The filtrate was collected and centrifuged to pellet cells, followed by resuspension in appropriate downstream culture or assay medium. Cell concentration and viability were assessed using a Countess II automated cell counter (Thermo Fisher Scientific) with Trypan Blue exclusion.

### Tumor-infiltrating lymphocyte (TIL) isolation and expansion

Patient-matched TILs were isolated by seeding 0.5 × 10^6^ dissociated patient tumor cells per well of a 24-well plate in ImmunoCult™-XF T Cell Expansion Medium supplemented with 6000 IU/mL recombinant human IL-2 (Miltenyi Biotec, 130–097-743). Cells were cultured for 7 days to promote initial TIL outgrowth. Subsequently, cultures were split and further expanded in fresh medium supplemented with a CD3/CD28/CD2 T cell activator (StemCell Technologies, 10971) to enhance T cell proliferation. Expanded TIL populations were then harvested, counted, and cryopreserved in CryoStor® CS10 (StemCell Technologies) for long-term storage in liquid nitrogen until use.

### Lung tumor organoid generation

Organoids were generated by encapsulating ∼ 2 × 10^5^ cells in 50 μL of Cultrex reduced growth factor basement membrane extract type 2 (“Matrigel”; R&D Systems, 3533-05-02,) and cultured with medium changes every 3 days [48]. The LTOs were maintained in basal medium supplemented with a cocktail of cytokines and growth factors. The basal medium is composed of Advanced DMEM/F12 (Thermo Fisher, 12634010), 1% GlutaMax (35050061), 1% HEPES (15630106), and 1% Penicillin-streptomycin (15140122). The complete LTO medium was prepared by supplementing the basal medium with 500 ng/mL R-Spondin 1 (PeproTech, 120-38), 25 ng/mL FGF-7 (PeproTech, 100-19), 100 ng/mL FGF-10 (PeproTech, 100-26), 100 ng/mL Noggin (PeproTech, 120-10-20UG), 500 nM A83-01 (R&D Systems, 2939/10), 1X B27 Supplement (Thermo Fisher, 12587010), 500 nM SB202190 (Sigma, S7067-5MG), 1.25 mM N-Acetylcysteine (Sigma, A9165-5G), 5 mM Nicotinamide (Sigma, N0636-100G), and 100 μg/mL Primocin (Invivogen, ant-pm-1). LTOs were cryopreserved in organoid medium supplemented with 10% FBS and 10% DMSO. For subsequent experiments, LTOs were thawed and resuspended in Matrigel. Droplets of 25 μL of the cell-suspended Matrigel solution were plated per well in a 24-well plate and cultured in organoid medium with medium changes every 3 days as described above.

### Lung tumor organoid characterization

Organoids were recovered from Matrigel using Cell Recovery Solution (Fisher Scientific, 08-774-405), fixed with 4% paraformaldehyde (PFA) for 15 min, washed with PBS, and resuspended in distilled water. The organoids were then spread onto glass slides (Leica, 3800200) and allowed to adhere by air drying. The organoids were then rehydrated in PBS and permeabilized with 0.1% Triton X-100 (Sigma-Aldrich, X100) in PBS for 15 min, followed by blocking with 1% BSA and 2.5% normal donkey serum in PBS (blocking buffer) for 1 hour. Samples were incubated overnight at 4 °C with primary antibodies against EpCAM (1:100, BioLegend, 324201), CK7 (1:100, Invitrogen, MA5-32173), and Napsin A (1:200, Abcam, ab166619) diluted in blocking buffer. After washing with PBS, samples were incubated with secondary antibodies, either Alexa Fluor 647 donkey anti-mouse (1:500, Thermo Fisher, A-31571) or Alexa Fluor 647 donkey anti-rabbit (1:500, Jackson ImmunoResearch, 711-605-152) in blocking buffer, along with Hoechst 33342 (2 μg/mL; Thermo Fisher, 62249), for 1 hour at room temperature. Samples were washed with PBS, mounted using mounting medium (Thermo Fisher, P36984), and imaged using an Andor Dragonfly Spinning Disk confocal microscope.

For hematoxylin and eosin (H&E) staining, air-dried organoids were first stained with Mayer’s hematoxylin (Electron Microscopy Sciences, 26043-06) for 3 min to visualize nuclei. Samples were rinsed with multiple changes of distilled water to remove excess hematoxylin. Nuclear bluing was achieved by incubating the samples in 1× PBS for 3 min, followed by washing with distilled water. Sections were then briefly rinsed in 95% ethanol for 1 min and counterstained with eosin (Electron Microscopy Sciences, 26051-21) for 1 min to visualize cytoplasm. The samples were then dehydrated through sequential 1 min washes in 95% ethanol (3X) followed by 100% ethanol (3X), and subsequently cleared in xylene (3X, 1 min each). Finally, samples were mounted in mounting medium and covered with a coverslip. The samples were imaged using a Keyence BZ-X710 microscope.

### Immunofluorescence of patient-biopsied tumor sections

For CD3 staining, Paraffin-embedded tissue sections were deparaffinized in xylene and rehydrated through a graded ethanol series. Antigen retrieval was performed in citrate buffer (Sigma-Aldrich, C999) for 15 minutes at 95 °C. Sections were then blocked in a solution containing 1% bovine serum albumin (BSA), 4% normal donkey serum, and 0.1% Triton X-100 in PBS for 1 hour at room temperature. Following blocking, sections were incubated overnight at 4 °C with a primary antibody against human CD3 (1:200 dilution, Novus Biologicals, NB600-1441SS) diluted in blocking buffer. After primary antibody incubation, sections were washed three times with PBS and then incubated with Alexa Fluor 647–conjugated donkey anti-rabbit secondary antibody (1:500 dilution, Jackson ImmunoResearch, 711-605-152) in blocking buffer for 1 hour at room temperature. Hoechst 33342 (2 μg/mL; Thermo Fisher, 62249) was included during secondary antibody incubation for nuclear staining. Sections were then washed with PBS and mounted using an appropriate mounting medium. Images were acquired using a Keyence microscope.

Quantitation of number of CD3^+^ cells in the patient tumor sections was performed using a custom CellProfiler pipeline. Briefly, DAPI^+^ cells and CD3^+^ cells were identified individually after background subtraction. Co-localization of CD3+/DAPI+ double-positive cells were determined, and the ratio of double-positive cells to total DAPI+ nuclei was quantified.

### Microvascular network formation

HUVECs were suspended in EGM-2MV medium supplemented with 4 U/mL thrombin (Sigma, T6634-500UN) at a concentration of 35 × 10^6^ cells/mL and kept on ice until use. NHLFs were suspended in EGM-2MV medium containing 4 U/mL thrombin at a concentration of 15 × 10^6^ cells/mL and kept on ice until use. Each cell suspension was separately mixed with an equal volume of 5 mg/mL fibrinogen (Thermo Fisher, J63267-03) in saline and injected into the microfluidic device: 15 μL of the HUVECs suspended in fibrin solution was injected into the central chamber *via* the access port, and 5 μL of the NHLF mixture was introduced into the fibroblast chamber *via* its inlet, yielding a ∼ 7:1 HUVEC:NHLF ratio. Care was taken to prevent pre-mature gelation of fibrin. The devices were transferred to the cell culture incubator, and cell-laden fibrin gels were polymerized at 37 °C for 25 min. EGM-2MV medium was then perfused through the side channels *via* media reservoirs and replaced daily for three days to allow formation of a stable microvascular network. The cell-loaded devices were monitored daily using phase contrast microscopy and fluorescence microscopy when fluorescently labeled cells were used. Images were acquired and visually confirmed the development of microvascular network, as indicated by the formation of interconnected tubular structures within the fibrin gel. The visual inspection also ensured maintenance of the fibrin gels’ structural integrity throughout the culture. The cell seeding and culture conditions were optimized to achieve stable microvascular network formation within three days. All cells were used between passages 3 and 5 and passage number was matched across all cell types within each experiment. Immunofluorescent staining of the microvascular networks, followed by quantitative image analysis was used to determine key vascular metrics, as described in detail below. Vascular perfusability and permeability were assessed by flowing the networks with medium containing fluorescent dextran, as detailed below.

### Microvascular network characterization

Microvascular network characterization using immunofluorescent staining was performed longitudinally starting on day 3 post-seeding. Day 3 was chosen based on longitudinal visualization and analysis of bright-field images of microvascular network formation within the device. Devices were fixed with 4% paraformaldehyde for 30 minutes, washed in 1X PBS, and permeabilized in PBS containing 0.1% Triton-X and blocked with PBS containing 0.1% Triton-X, 1.5% bovine serum albumin, and 2.5% normal donkey serum for 2 hours at 25 °C. Primary antibodies (1:200, VE-cadherin; R&D, AF938) diluted in blocking buffer were perfused into the devices and incubated overnight at 4°C. Devices were then washed with 1X PBS containing 0.1% Tween-20 (wash buffer). Alexa Fluor 647-conjugated donkey anti-goat (1:200, Invitrogen, A-21447) diluted in blocking buffer was introduced into devices and incubated overnight at 4 °C. Devices were subsequently washed in wash buffer and imaged using a Keyence BZ-X710 fluorescence microscope or Andor Dragonfly Spinning Disk Confocal Microscope. Maximum intensity z-projections were generated from z-stack images and analyzed using AngioTool v0.6a [26]. AngioTool allows semi-automated quantification of vascular network parameters such as average vessel length, lacunarity, and junction density. Images were processed according to the NCI AngioTool user guide (https://ccrod.cancer.gov/confluence/display/ROB2/Quick+Guide). All additional image analyses were performed using ImageJ (v2.14.0). A minimum of three microvascular network ROIs per device were analyzed across at least four independent devices.

To measure vascular permeability, 10 μL of 2000kDa FITC-Dextran (Sigma, FD2000S) was added to the designated media inlet reservoir to generate a pressure-driven flow across the microvasculature. Time-lapse fluorescent images were acquired every 30 s for 5 min. In ImageJ, regions of interest were defined to include adjacent extravascular space and vessel segments with diameters <50 μm to ensure near-circular cross-sections. Fluorescence intensity was determined over time within each region. The permeability coefficient was calculated using the following equation:

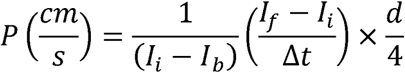

where *l_b_, l_i_* and *l_f_* represent the mean fluorescence intensity within the measurement window background, initial, and final time points, respectively. Δt denotes the time interval between the initial and final measurements (s) and *d* is the average vessel diameter within the measurement window.

### Lung tumor organoid (LTO) implantation

LTOs were implanted into the LTO chamber of the devices after three days of HUVEC and NHLF culture in the vLTO devices. Implantation into an established vascular bed was crucial, as the perfusable networks did not form when LTOs, HUVECs, and NHLFs were seeded simultaneously or when LTOs were seeded before day three of microvascular network culture. LTOs were generated and cultured independently in LTO medium as described above. The LTOs were collected from the culture and centrifuged at 100xg for 3 minutes. LTOs were released from the Matrigel by incubating in Cell Recovery Solution (Corning, 354253) at 4 °C for 15 min. For longitudinal imaging experiments, LTOs were stained with CellTracker Green CMFDA (Thermo Fisher, C7025) according to the manufacturer’s instructions. Briefly, the Matrigel-released LTOs were incubated with CellTracker Green CMFDA dye in PBS (10 μM) at 37 °C for 30 minutes. LTOs were then washed with LTO media and centrifuged at 100xg for 3 min. This step was repeated for at least two washes. LTOs were resuspended in a hydrogel mixture composed of 60% (v/v) Matrigel and 7 mg/mL fibrinogen, with a final fibrinogen concentration of 2.5 mg/mL. Thrombin was added to the LTO-matrix mixture to a final concentration of 2 U/mL, mixed, and 5 μL of the fibrin gel precursor was immediately introduced into the organoid compartment. Gelation was allowed to proceed for 30 min at 37 °C. After ensuring complete gelation, a 1:1 mixture of LTO:EGM-2MV medium was added to media inlet ports. A 15 μL volume of 50 ng/mL VEGF (Thermo Fisher, 100-20-10UG) dissolved in the same medium was placed on top of the organoid chamber and replenished during media changes to establish a chemokine gradient and promote anastomosis between the vascular network and the tumor organoid. Additional care was taken to prevent medium evaporation; the devices were placed in a humidified chamber and subsequently maintained in a cell culture incubator at 37 °C with 5% CO₂ and 21% O₂ with daily media change before performing experiments and analyses.

### Characterization of vascularized LTO-on-chip devices

Vascularized LTO-on-chip devices were fixed and stained as described above. The LTOs and vasculature were stained with the primary antibody (anti-human EpCAM; Biolegend, 324201 and anti-human VE-Cadherin; R&D Systems, AF938, respectively), which was introduced into the LTO chamber and incubated overnight at 4 °C. Devices were then washed with wash buffer, and incubated with the secondary antibody (Alexa Fluor 488 AffiniPure Donkey Anti-Mouse IgG (H+L), Jackson ImmunoResearch, 715-545-150 and Alexa Fluor 594 AffiniPure Donkey Anti-Goat IgG (H+L), Jackson ImmunoResearch, 705-585-147) overnight at 4 °C. The devices washed with wash buffer were imaged using a Keyence BZ-X710 Fluorescence Microscope or Andor Dragonfly Spinning Disk Confocal Microscope. Approximately 3-4 LTOs per device and multiple devices fabricated and cultured independently were imaged. Confocal images (IMS files) were imported into Imaris software, and surface reconstructions were generated for both the microvasculature and LTOs. A distance transformation analysis was performed to quantify the distance between the microvasculature and LTO surfaces. Surface-surface co-localization analysis was used to quantify the contact area between the LTOs and the microvasculature.

Vascularized devices with and without the LTO incorporation were used to determine the effect of LTOs on the microvascular network formation. Similarly, the effect of a perfusable microvascular network on organoid growth, LTOs cultured within perfusable microvascular networks were compared with LTOs maintained outside the device under static conditions. For these comparisons, organoids from the same culture batch were used and maintained under otherwise identical media and incubation conditions. Organoids were monitored over a period of four days. Phase-contrast images were acquired daily using an inverted microscope under consistent imaging settings. Organoid growth was quantified by measuring projected area using ImageJ (NIH). For each condition, regions of interest were manually defined, and the two-dimensional area of individual organoids was calculated and normalized to the initial area at day 0 to account for size variability at seeding.

### Chemotherapeutic response in vascularized LTOs and conventional LTO cultures

Cisplatin (Sigma, 232120) was diluted to a final concentration range of 0 –100 μM in a 1:1 mixture of LTO medium and EGM-2MV medium. The drug-containing medium was introduced into the media inlet reservoir of the device, enabling perfusion across the microvascular network and into the tumor organoid compartment by pressure driven flow. Devices were maintained under continuous perfusion for 24 hours at 37 °C and 5% CO₂. Following cisplatin exposure, propidium iodide (PI; 1 μg /mL) was perfused through the device for 30 min at 37 °C in the dark. Organoids were then imaged using a Keyence BZ-X710 fluorescence microscope under consistent acquisition settings across conditions. Quantification of cell death was performed using ImageJ (NIH). Briefly, PI channel (red fluorescence) images were imported, and masks delineating individual LTOs within the organoid compartment were manually generated. Mean fluorescence intensity within each mask was measured and normalized to the corresponding mask area to account for organoid size differences. For comparison in static culture, LTOs were encapsulated in 70% (v/v) Matrigel at a density of 10,000 cells per 5 μL droplet and cultured in a 96-well plate under standard culture conditions. Cisplatin was prepared as described above and added directly to the culture medium. After 24 h of treatment, cell viability was assessed using the CellTiter-Glo 3D Cell Viability Assay (Promega, G9682) according to the manufacturer’s instructions. Briefly, 150 μL of CellTiter-Glo 3D reagent was added to each well, and contents were mixed thoroughly by pipetting at least 10 times to ensure complete lysis. Plates were equilibrated at room temperature for 25 min to stabilize the luminescent signal, which was subsequently measured using a Tecan Infinite 200 Pro plate reader with an integration time of 1 s. Luminescence values were normalized to untreated controls to determine relative viability

### CD8^+^ T cell infiltration

The vascularized LTO were cultured for 7 days prior to immune cell perfusion experiments. THP-1 monocytes and primary CD8^+^ T cells were labeled with fluorescent trackers by incubation in 500 nM CellTracker Deep Red (Thermo Fisher, C34565) and 1 μM CellTracker Red CMTPX (Thermo Fisher, C34552), respectively, for 20 min at 37 °C in the dark. Following staining, cells were washed at least twice with their respective culture media and centrifuged at 125 × g for 3 min between washes to remove excess dye. Approximately 1 × 10L THP-1 monocytes suspended in a 1:1 mixture of LTO medium and THP-1 medium were introduced into the vascular channel of the device and perfused under pressure-driven flow. Cells were allowed to adhere to the microvascular network for 2 h at 37 °C and 5% CO₂, after which non-adherent cells were removed by perfusion with fresh medium. Subsequently, ∼5 × 10L CD8⁺ T cells suspended in a 1:1 mixture of LTO medium and T cell culture medium were introduced into the vascular channel. Devices were maintained under static conditions for 2 h to facilitate T cell adhesion to the endothelium, after which perfusion was resumed. Following immune cell introduction, devices were cultured under continuous perfusion for 24 h to allow T cell trafficking and infiltration into the organoids. To assess the role of monocytes in T cell recruitment, parallel conditions were established with and without the incorporation of THP-1 cells. After 24 h, devices were imaged using an Andor Dragonfly Spinning Disk Confocal Microscope. At least two vLTOs per device were analyzed across multiple independently fabricated and cultured devices. Confocal image stacks (IMS format) were imported into Imaris for analysis. Three-dimensional surface reconstructions were generated for both CD8⁺ T cells and LTO structures. A distance transformation analysis was performed to quantify the spatial relationship between T cells and organoids. For quantitative comparison, the number of CD8⁺ T cells located within 100 μm of the LTO surface was determined.

### Generation of β2-microglobulin-knockout HUVECs

Low passage HUVECs were suspended in P5 Primary Cell 4D-Nucleofector Solution and electroporated with Cas9 ribonucleoprotein (RNP) complexes using a 1:2 molar ratio of Cas9 to guide RNA. Electroporation was performed using a Lonza 4D-Nucleofector® X Unit, according to the manufacturer’s instructions for the P5 Primary Cell Nucleofector® X Kit L (V4XP-5024). Two single guide RNAs (sgRNAs) targeting distinct regions within the *B2M* locus were designed and evaluated to identify the most efficient editing site. Cas9 protein (TrueCut Cas9 v2; Thermo Fisher, A36499) and synthetic sgRNAs (TrueGuide sgRNA; Thermo Fisher, A35514) were assembled into RNP complexes immediately prior to electroporation following the manufacturer’s recommendations. Following electroporation, HUVECs were cultured for 9 d to allow for gene editing and protein turnover. Cells were then collected and stained with an anti-β2-microglobulin (B2M) antibody conjugated to Alexa Fluor 647 (MA5-18119; Thermo Fisher) for 30 min at 4 °C in the dark. B2M-deficient (*B2M*⁻/⁻) cells were isolated by fluorescence-activated cell sorting using a Beckman Coulter MoFlo Astrios EQ. Sorted cells were re-plated, expanded under standard endothelial growth conditions, and subsequently cryopreserved for downstream experiments.

### Assessment of T cell activation

Changes in the ability of β2M-knockout (β2M-KO) HUVECs to activate T cells were evaluated relative to unedited control HUVECs by quantifying interferon-γ (IFNγ) secretion as a functional readout. β2M-KO or unedited HUVECs were seeded in 96-well plates and cultured using EGM-2MV medium until ∼90% confluence. Primary CD8⁺ T cells were then added to HUVEC monolayers at an effector-to-target (T cell:HUVEC) ratio of 20:1 and co-cultured in T cell medium for 24 h at 37 °C and 5% CO₂. Following incubation, the medium was collected and IFNγ levels were quantified using a human IFNγ ELISA kit (R&D Systems, DIF50C) according to the manufacturer’s instructions. Absorbance was measured using a microplate reader, and cytokine concentrations were calculated by interpolation from standard curves generated using a four-parameter logistic (4PL) fit in GraphPad Prism (v9.5.1). A minimum of 3-4 samples per experimental group were used and results were normalized to control HUVEC–T cell co-cultures.

### Microvascular network formation from *B2M* knockout HUVECs

Microvascular networks were formed by β2M-KO HUVECs similar to unedited HUVECs as described earlier and characterized.

### Patient-matched TIL infiltration

Microvascular networks were formed using β2M-KO HUVECs and LTOs were transplanted into the designated compartment and cultured in a 1:1 mixed medium of LTO and EGM2-MV medium as described above. The vascularized LTO was then exposed to THP-1 monocytes and non-adherent cells were subsequently removed by perfusion of fresh medium through the device similar to that described earlier. Approximately 50,000 patient-matched TILs, labeled with CellTracker Red CMTPX following the same protocol used for CD8⁺ T cells and resuspended in a 1:1 mixture of LTO medium and EGM-2MV medium, supplemented with either vehicle control or 10 μg mL⁻¹ anti-PD-1 antibody (BioXcell, SIM0003). TILs were introduced into the vascular channel and allowed to circulate through the microvascular network to enable interaction with LTOs, followed by 24 h of culture under continuous perfusion, as described above. TIL infiltration into LTOs was assessed by imaging using an Andor Dragonfly Spinning Disk Confocal Microscope. At least two vascularized LTOs per device were analyzed across multiple independently fabricated and cultured devices. Confocal image stacks (IMS format) were imported into Imaris, where three-dimensional surface reconstructions were generated for both TILs and LTOs. A distance transformation analysis was performed to quantify spatial relationships between TILs and LTO surfaces. For quantitative assessment, the number of TILs located within 100 μm of the LTO surface was determined.

Experiments determining whether targeting hyperangiogenic signaling could improve T cell recruitment evaluated concentration-dependent effects of vascular endothelial growth factor (VEGF) antibody (BioXcel, SIM0007) on promoting T cell infiltration into patient-derived LTOs, using LTOs from patient 2 and corresponding autologous TILs. The experiments were performed using the same vascularized LTO platform and immune perfusion workflow described above. Briefly, vascularized LTOs were perfused with 1:1 LTO:EGM2-MV medium supplemented with 10 μg/ml of Anti-PD1 and increasing concentrations of anti-VEGF antibody (0, 10, or 100 μg mL⁻¹; BioXcell, SIM0007). TILs were then perfused through the vascular network and allowed to interact with the LTO compartment for 24 h under continuous perfusion as described above. TIL infiltration was subsequently quantified as described above.

### Sample preparation for single-cell RNA sequencing

LTOs along with THP-1 and matched TILs were retrieved by enzymatic digestion using a collagenase/dispase solution in PBS with a final concentration of 0.1 mg/mL collagenase and 0.5U/mL of dispase (Worthington 9001-12-1, STEMCELL Technologies cat. no. 07913) at 37 °C or 15 min with gentle agitation. The reaction was quenched by resuspending the sample in 10 mL of PBS, followed by centrifugation at 500 × g for 5 min. Cell pellets were resuspended in 400 μL TrypLE Express (Gibco, 12604013) per sample, and gently dissociated by pipetting, followed by incubation at 37 °C for 20 min to ensure complete single cell dissociation. The reaction was quenched by addition of 5 mL basal medium and centrifugation at 500xg for 5 min with low brake. Cell pellets were resuspended in 1 mL FACS buffer (PBS supplemented with 2 mM EDTA, 5% FBS, and 0.1% non-essential amino acids). Approximately 1 μL of propidium iodide (PI; ThermoFisher, P1304MP) was added (1 μL per 900 μL cell suspension) immediately prior to sorting. Live, PI-negative singlet cells were sorted using a Beckman Coulter MoFlo Astrios EQ High Speed Sorter and collected into low-binding tubes containing 100 μL chilled FACS buffer. Sorted cells were adjusted to a final cell concentration of 8,000 cells/μL to ensure encapsulation of single cells using the Chromium Next GEM Single Cell 3’ Reagent Kits v3.1 (Dual Index, 10x Genomics).

### Single-cell RNA sequencing library preparation

Single-cell RNA sequencing libraries were prepared using the Chromium Next GEM Single Cell 3’ Reagent Kits v3.1 (Dual Index; 10x Genomics, PN-1000268) according to manufacturer’s instruction. In brief, ∼18,000 single cells per sample were loaded onto a Chromium Next GEM Chip G (PN-1000120) and processed using a Chromium X Controller for co-encapsulation with barcoded beads at a target capture rate of 10,000 individual cells per sample. Gel Bead-in-Emulsions (GEMs) were generated, followed by reverse transcription, emulsion breaking, and cDNA recovery using Dynabeads MyOne SILANE beads (ThermoFisher, PN-2000048). cDNA was amplified and purified using SPRIselect beads (Beckman Coulter PN-B23318). Dual-indexed 3’ gene expression libraries were constructed through enzymatic fragmentation, end-repair, A-tailing, adapter ligation, index PCR, and double-sided size selection. Final libraries were quantified using an Agilent TapeStation and sequenced on an Illumina NovaSeq X Plus system using a 10B flow cells, generating ∼ 300 million paired reads per sample (28 bp Read 1, 10 bp i7 index, 10 bp i5 index, and 90 bp Read 2).

### Computational analysis

Raw sequencing data were demultiplexed FASTQ files and processed using Cell Ranger v6.1.2 (10x Genomics) and aligned to the GRCh38 reference genome to generate gene-barcode matrices. Doublets were identified and removed using the Solo algorithm implemented within the scVI variational autoencoder framework [49]. Cells with fewer than 200 detected genes, more than 10,000 detected genes, or with mitochondrial transcript count exceeding 15% were excluded.

Downstream analysis was performed in Seurat v5.3.0. Data were log-normalized using NormalizeData, and variable features were identified using FindVariableFeatures. For dataset integration, 3,000 shared highly variable genes were selected, and integration anchors were identified using FindIntegrationAnchors with 30 principal components. Batch correction and data integration were performed using IntegrateData. Dimensionality reduction was performed using principal component analysis (PCA), followed by Uniform Manifold Approximation and Projection (UMAP) visualization using the first 30 principal components. A shared nearest-neighbor (SNN) graph was constructed, and unsupervised clustering was performed using the Louvain algorithm.

Cluster-specific marker genes were identified using FindAllMarkers (min.pct = 0.25, logfc.threshold = 0.25, only.pos = TRUE). Cell types were manually annotated using published datasets and CellMarker2.0 [50]. Differentially expressed genes (DEGs) were identified using FindMarkers (p_val_adj < 0.05, |avg_log2FC| > 1.5). Volcano plots were generated using the EnhancedVolcano R package. The top 100 DEGs in each direction (ranked by adjusted p-value) were used as input to identify enriched gene ontologies using EnrichR against GO Biological Process 2025. The top 20 enriched terms were plotted by Odds Ratio and rank ordered by adjusted p-value (p_val_adj < 0.05). Code used for single-cell RNA-seq analysis and visualization was developed with assistance from large language models and manually verified by the authors.

## SUPPLEMENTARY MATERIAL

**Supplementary Table S1:** NSCLC Patient Demographics and Clinical Diagnosis.

| ID | Age | Gender | Race | Histology | Grade | Primary Site | Metastatic Site | Pathologist Comments |
| --- | --- | --- | --- | --- | --- | --- | --- | --- |
| 1 | 62 | M | Caucasian | Invasive mucinous adenocarcinoma | Moderately differentiated | Lung | None | < 1 lymphocyte per 10 tumor cells |
| 2 | 63 | F | Caucasian | Invasive mucinous adenocarcinoma | Moderately differentiated | Lung | None | ~5 IEL's per 10 tumor cells |

**Supplementary Figure S1: Formation of microvascular networks and morphological evolution of lung tumor organoids (LTOs).** (a) Schematic diagram of the device with dimensions; R2 = 4 mm, Ø3 = 3 mm. (b) Step-wise timeline illustrating the experimental workflow for establishing a vascularized LTO-on-a-chip from day 0 to 7. (c) Bright-field and fluorescence images showing HUVEC network formation in the central chamber of the device. (d) Phase contrast images of LTO in a 96-well plate and vLTO in the device after seeding at 0 hr and 96 hr.

**Supplementary Figure S2: 3D analysis of vasculature and LTO distances.** Rendering confocal Z-stack images into 3D surfaces of vasculature (red) and LTO (green) using IMARIS software, followed by applying a distance transformation and overlaying the surface image with the distance-transformed image.

**Supplementary Figure S3: Cytotoxicity of cisplatin on HUVEC monolayer.** Fluorescence images and quantification of dead cells in HUVEC monolayers cultured in 96-well plates treated with varying concentrations of cisplatin; N = 3 wells, One-way ANOVA with Tukey’s post-hoc multiple comparisons testing; ** = p-value < 0.05.

**Supplementary Figure S4: Validation of *B2M* knockout in HUVECs.** (a) Flow cytometry analysis of *B2M* expression in wild-type (Control) and *B2M* knockout (KO) HUVECs. (b) TIDE analysis of genomic DNA from *B2M*-KO HUVECs with the distribution of indels (insertions and deletions) at the CRISPR target site. (c) Sanger sequencing of the *B2M*-KO region confirming mutations at the guide RNA target site compared to the wild-type (WT) sequence.

**Supplementary Figure S5: Microvascular network formation with *B2M*-KO HUVECs and reduced CD8^+^ T cell activation in co-culture with *B2M*-KO HUVECs.** (a) Fluorescence microscopy image of *B2M*-KO HUVECs and corresponding key morphological parameters compared with wild-type (WT) HUVECs; N = 3 devices per group. (b) ELISA analysis of IFN-γ secretion in co-cultures of CD8^+^ T cells with *B2M*-KO HUVECs and WT HUVECs.

**Supplementary Figure S6: Patient tumor baseline characteristics.** (a) Representative H&E stains of Patient 1 biopsied tumor section and adjacent normal tissue section. (b) Representative H&E stains of Patient 2 biopsied tumor section and adjacent normal tissue section. (c) Fluorescence microscopy image of Patient 1 and Patient 2 biopsied tumor section immunostained for CD3 (red) and DAPI (gray). (d) Quantification of CD3^+^ cells as a percentage of total (DAPI^+^) cells.

**Supplementary Figure S7: Single-cell RNA sequencing cell annotation and enrichment.** (a-d) UMAP visualization colored by canonical cell marker expression. (e-j) Differentially expressed genes for Regulatory T cells, Cycling T cells, Monocytes, NK cells, Naïve central memory T cells, and Epithelial cells respectively (|Log2 fold change| ≥ 1.5, adjusted p value < 0.05).

## ACKNOWLEDGEMENTS

This study is partially supported by funding from NCI/NIH (CA251407). This work was performed in part at the Duke University Shared Materials Instrumentation Facility (SMIF), a member of the North Carolina Research Triangle Nanotechnology Network (RTNN), which is supported by the National Science Foundation (Award No. ECCS-2025064) as part of the National Nanotechnology Coordinated Infrastructure (NNCI). We gratefully acknowledge the Duke BioRepository and Precision Pathology Center, Duke Light Microscopy Core Facility, and the Duke Cancer Institute Flow Cytometry Shared Resource for their invaluable support and assistance in this work. We thank all members of the Varghese laboratory for insightful discussions and feedback.

## Contributions

NRN and SV conceptualized the study. NRN designed the studies and performed the experiments with help from SM and SMP. SM, RK and GC performed experiments and sample preparation related to single-cell RNA-sequencing. RK analyzed the single-cell RNA-sequencing data with feedback from ZJ. NRN generated CRISPR-Cas9-edited *B2M*-KO HUVECs with assistance from NA. NRN and SV wrote and edited the manuscript.

