## Supplementary Figures for "Patient-Specific Vascularized Lung Tumor Organoids for Tumor-Immune Profiling"

### Supplementary Figure S1

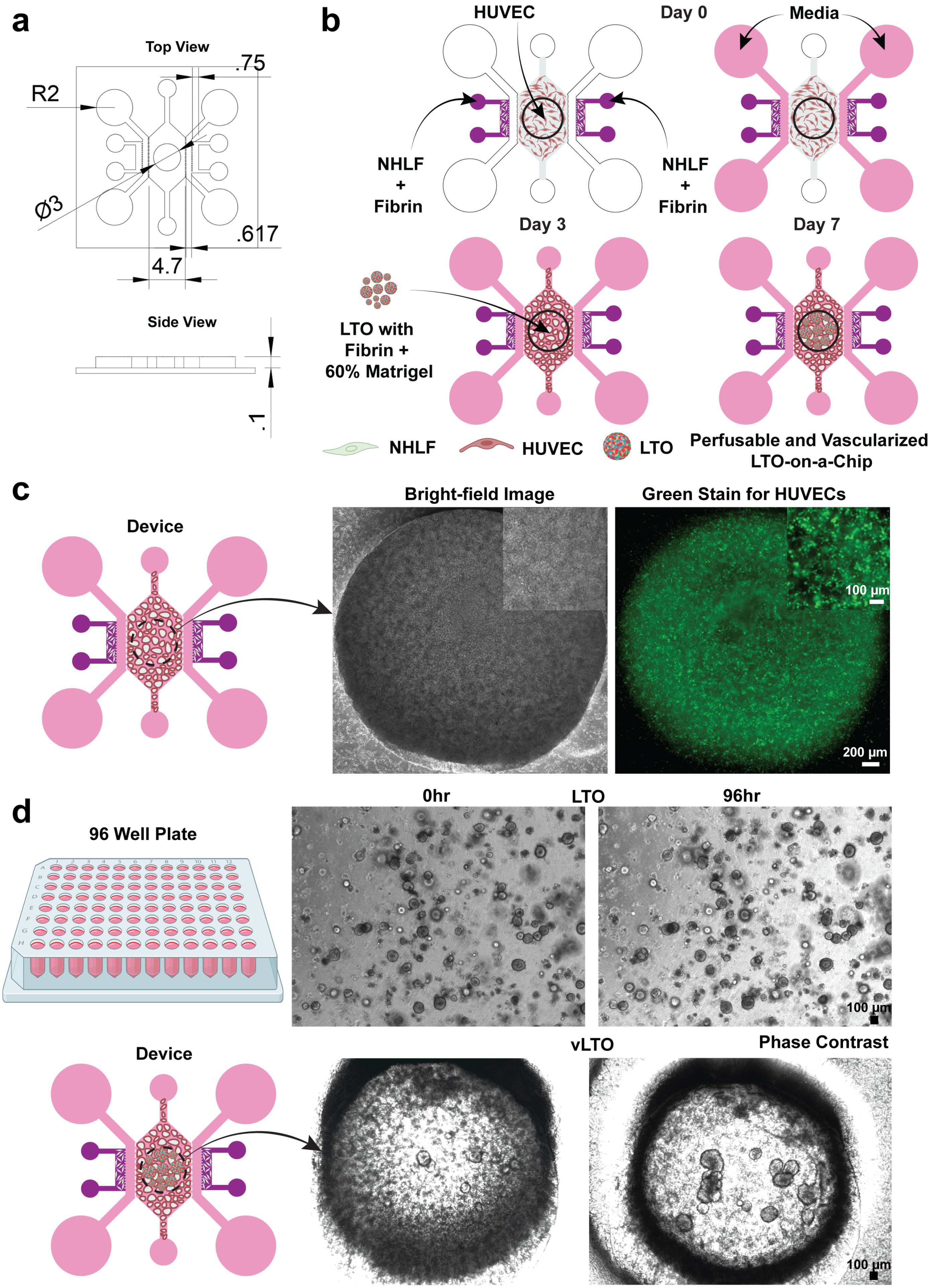

### Supplementary Figure S2

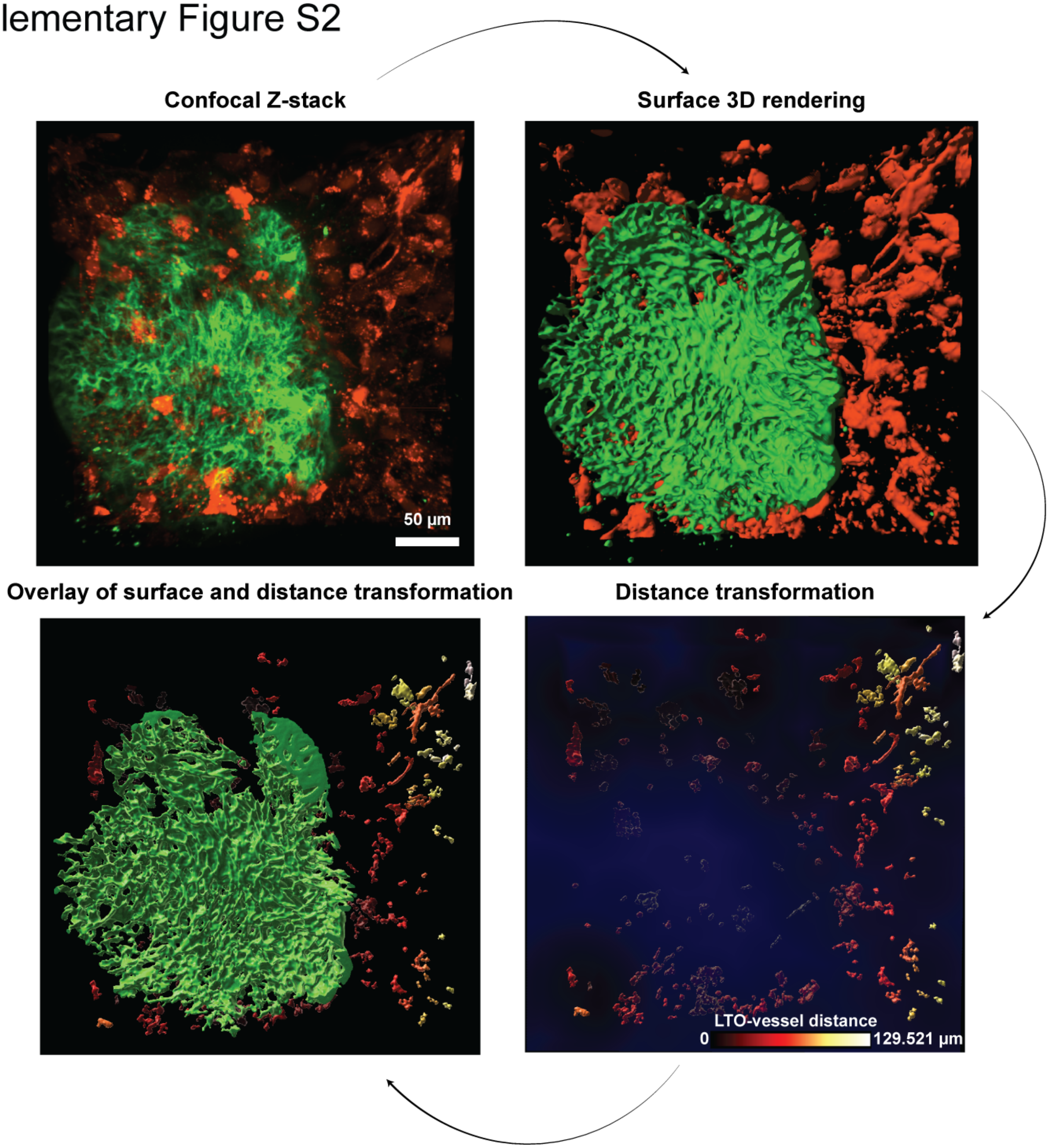

Supplementary Figure S3

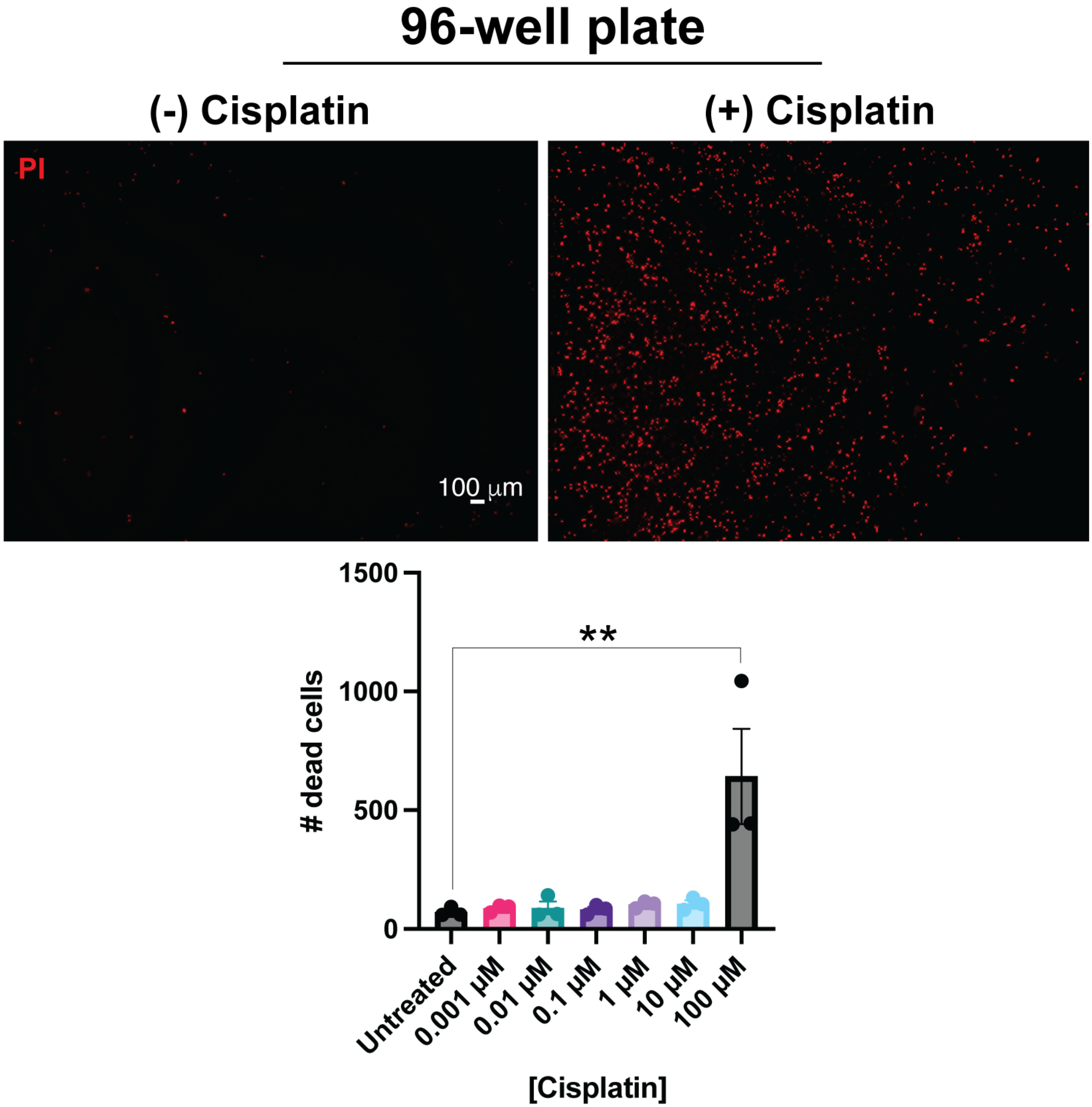

Supplementary Figure S4

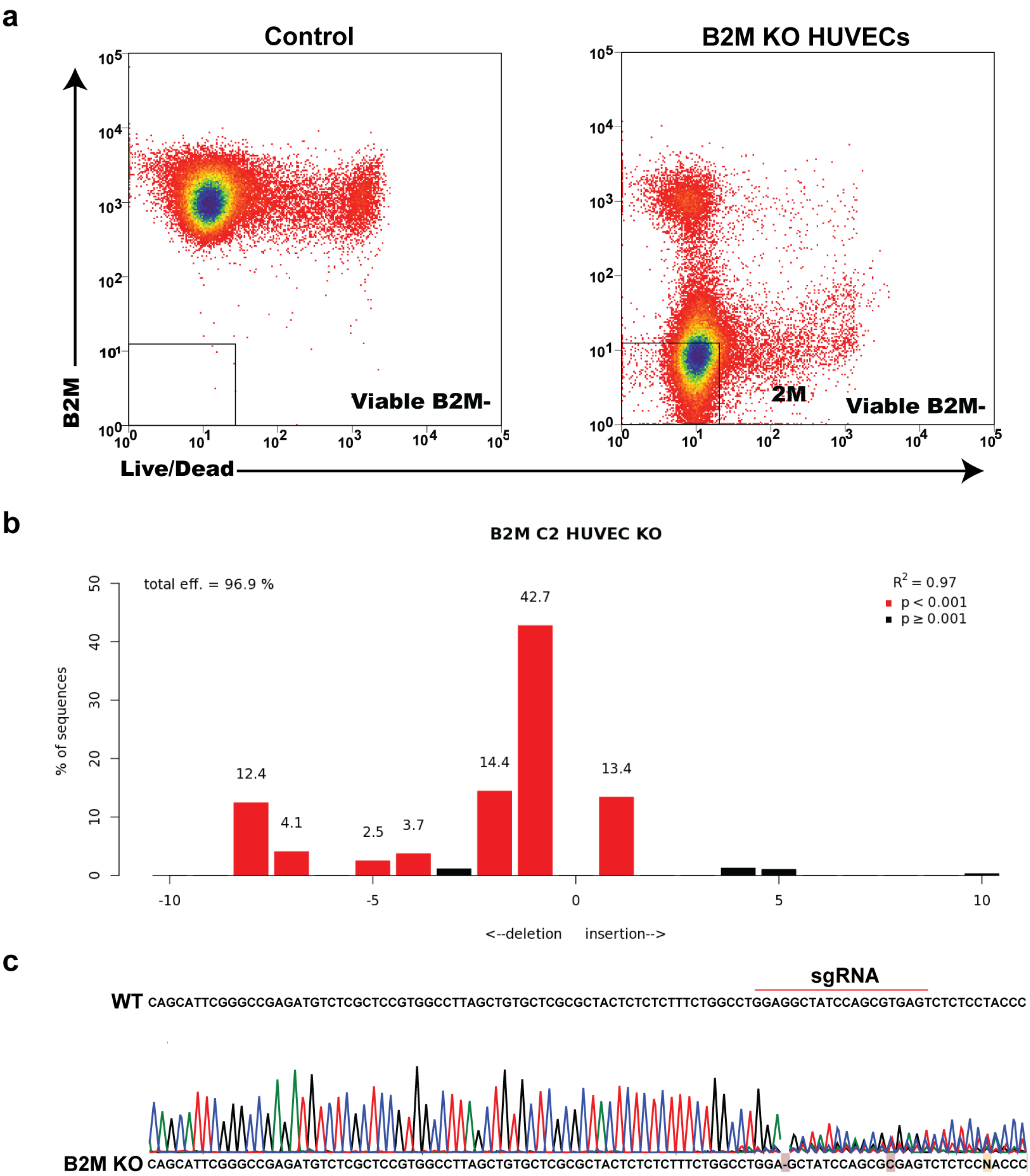

Supplementary Figure S5

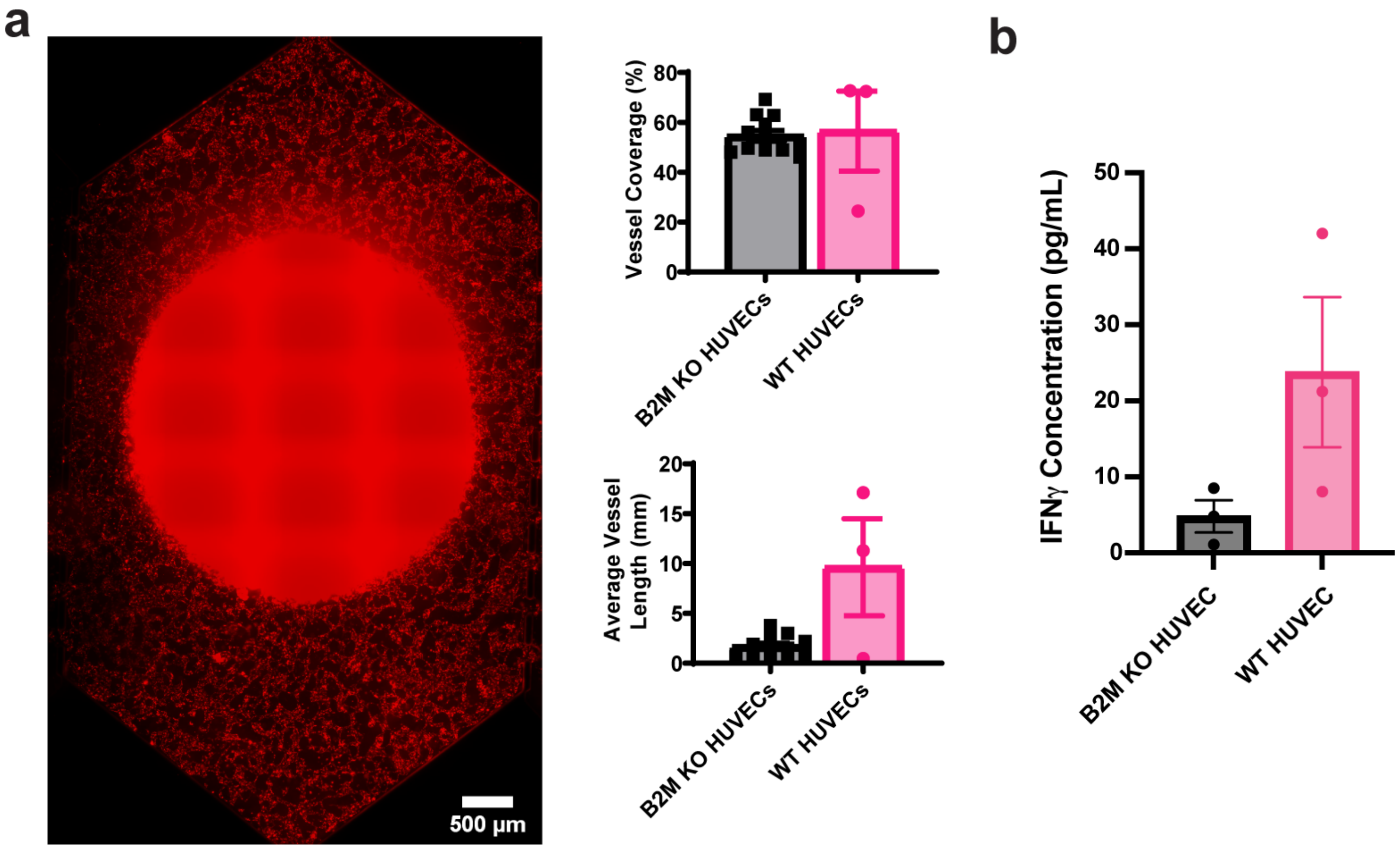

### Supplementary Figure S6

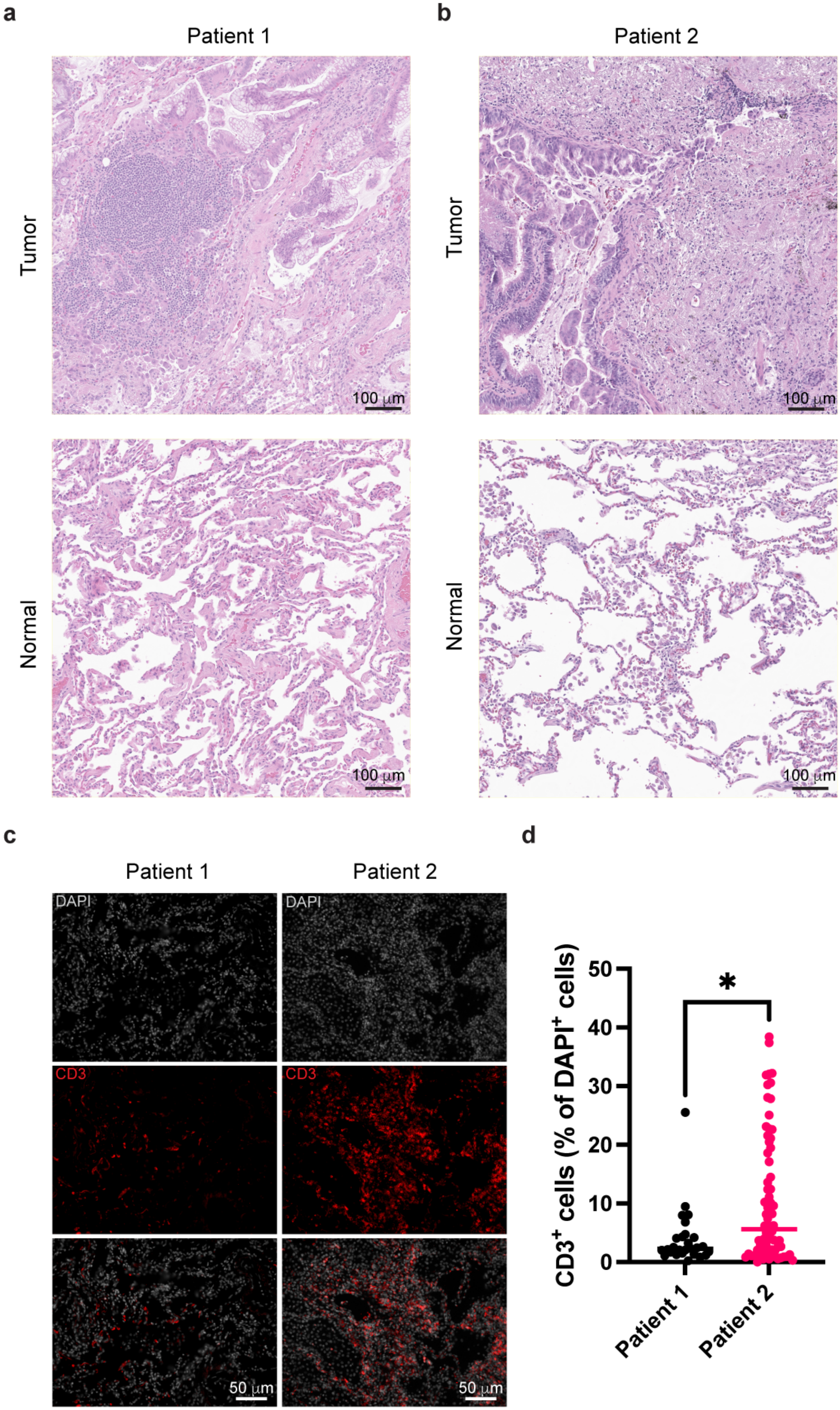

### Supplementary Figure S7

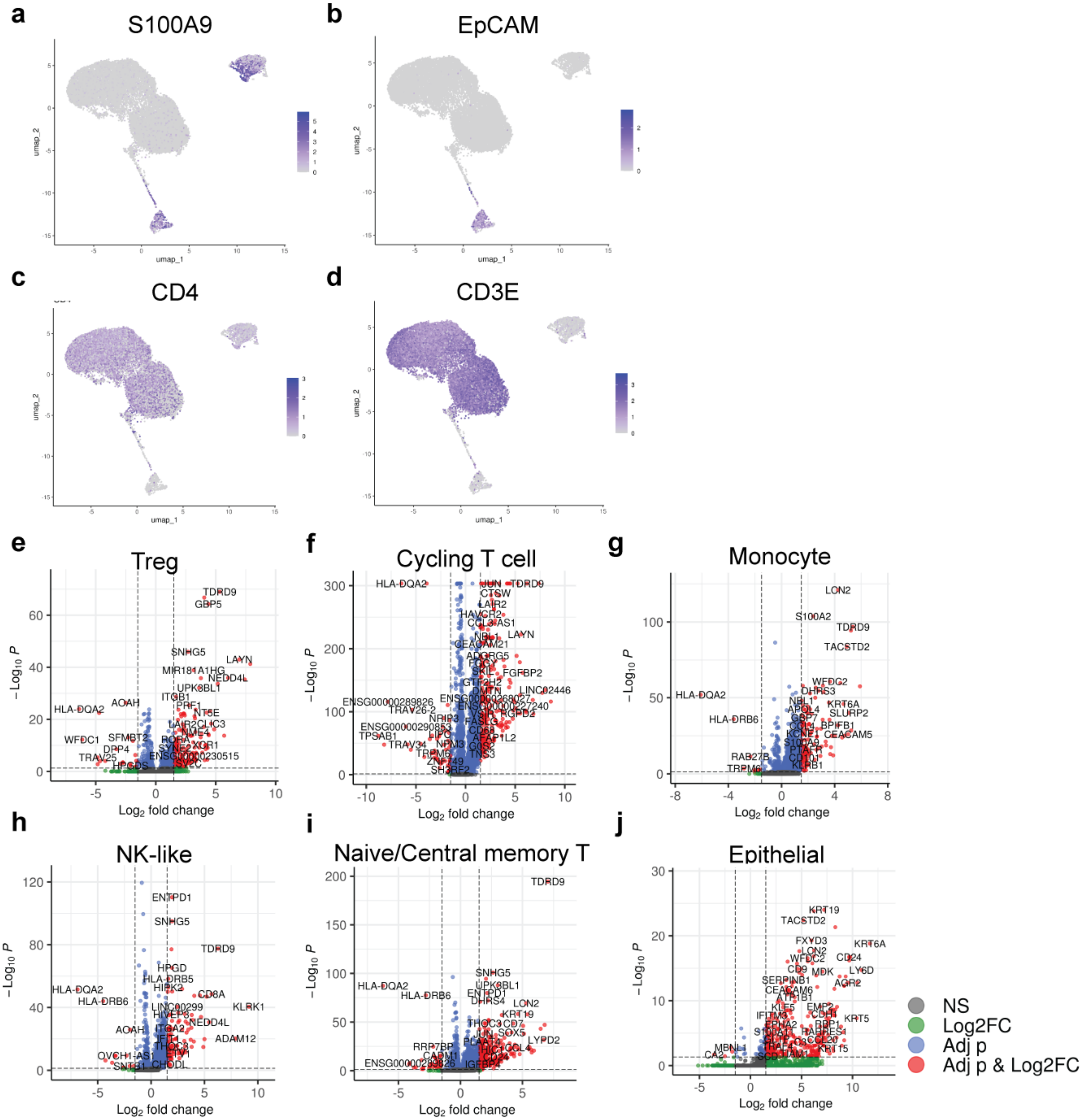
